# Single-cell and spatially resolved atlas of pancreatic cancer reveals immunophenotypes associated with clinical outcome

**DOI:** 10.1101/2025.02.08.637283

**Authors:** Gabriel F. Pozo de Mattos P., Marvin Paulo Lins, Julia Fontoura, Alessandro Bersch Osvaldt, Simone Marcia dos Santos Machado, Eduardo Cremonese Filippi-Chiela, Cristina Bonorino

**Affiliations:** Basic Health Sciences Department, Federal University of Health Sciences of Porto Alegre (UFCSPA), Immunotherapy Laboratory–UFCSPA, R. Sarmento Leite, 245-Centro Histórico, Porto Alegre, 90050-170, Brazil; Experimental Research Service, Hospital de Clínicas de Porto Alegre. Ramiro Barcelos 2350. 90035-903 Porto Alegre / RS-Brazil; Department of Morphological Sciences, Universidade Federal do Rio Grande do Sul. Sarmento Leite 500. 90035-903 Porto Alegre / RS-Brazil; Biotechnology Center, Universidade Federal do Rio Grande do Sul, Porto Alegre, RS, Brazil; Medical School, Universidade Federal do Rio Grande do Sul, Porto Alegre, RS, Brazil; Digestive Surgery Service, Biliary and Pancreas Group, Hospital de Clínicas de Porto Alegre, Porto Alegre, RS, Brazil; Graduate Program in Medicine: Surgical Sciences, Universidade Federal do Rio Grande do Sul, Porto Alegre, RS, Brazil; Pathology Service, Hospital de Clínicas de Porto Alegre, Porto Alegre, RS, Brazil

**Keywords:** single-cell, pancreatic cancer, spatial transcriptomics, immunophenotypes

## Abstract

Pancreatic ductal adenocarcinoma (PDAC), accounting for 90% of pancreatic neoplasms, is characterized by its poor prognosis, with a 5-year survival rate of only 12%. Most patients are diagnosed with metastatic or locally advanced disease, leaving only 15% eligible for curative resection. PDAC exhibits resistance to chemotherapy, targeted therapies, and immunotherapy, largely due to its highly heterogeneous tumor microenvironment (TME). In this study, we performed an integrative analysis of publicly available scRNA, spatial transcriptomics, and bulk RNA sequencing datasets to investigate the influence of TME composition and tumor architecture on PDAC progression, treatment response, and clinical outcomes. We identified TME subtypes with distinct cellular compositions, functional signatures, and immunomodulatory cell-cell interactions. Spatially distinct cellular niches and gene modules revealed heterogeneity across primary tumors and metastatic lesions. Deconvolution of these spatial niches in a large cohort of bulk RNA samples uncovered unique clusters associated with patient survival, providing novel insights into TME biology and its clinical implications. These findings underscore the importance of integrating multi-omics approaches to unravel the complexity of the PDAC TME and highlight its potential to inform therapeutic strategies and improve patient outcomes.

## Introduction

Pancreatic ductal adenocarcinoma (PDAC) represents 90% of pancreatic neoplasms (1). Unfortunately, most patients already present metastatic lesions at the moment of diagnosis and around 35% have locally advanced disease (2). Due to the late diagnosis, only 15% of the patients are eligible for resection, which is the only curative treatment available until today. PDAC is highly resistant to standard chemotherapy/chemoradiotherapy and remains refractory to targeted therapies. Despite all efforts to improve patient outcomes, the 5-year survival rate just reached 12% last year (3). These challenges arise from the highly heterogeneous tumor microenvironment (TME) observed in PDAC (4).

TME is a complex ecosystem formed by stromal cells, immune cells, and tumor cells beyond the extracellular matrix (ECM). Non-tumoral cells engage in heterotypic intercellular interactions among themselves and with malignant cells. TME composition is dynamic throughout carcinogenesis, from premalignant lesions to established metastasis (5–8). For instance, in many tumors, including PDAC, infiltration by competent immune cells is most frequently associated with response to therapy, improved survival, and considered to prevent tumor progression (9–15). This has not been the case for PDAC, where TME is characterized by the high prevalence of cancer-associated fibroblasts (CAFs) and myeloid cells, which sustain malignant cells and promote an immunosuppressive TME associated with few cytotoxic lymphocytes (4).

Single-cell RNA (scRNA) has provided valuable insights into the complexity of the TME in PDAC revealing extensive heterogeneity of malignant cells, CAFs, and immune milieu (7,8,10,11, 16–23). However, the clinical impact of single-cell studies remains limited due to the absence of spatial context (24). Recent reports unraveled the impact of spatial organization on tumor progression, therapy response, and prognosis in multiple cancers (25–28). Spatial transcriptomics (stRNA) emerged as a transformative tool to dissect complex cell-cell interactions within TME (29,30).

Here, we performed extensive data integration of publicly available scRNA, stRNA, and bulk RNA datasets to comprehend the influence of TME composition and tumor architecture in disease progression, therapy response, and clinical outcome. We identified cellular composition differences associated with distinct tissue of origin and treatment-associated changes. We uncovered TME subtypes with distinct composition, functional signatures, and cell-cell interactions involved with immunomodulation. By integrating scRNA with spatial transcriptomics, we were able to identify cellular niches and spatial gene modules uncovering spatial heterogeneity across primary tumors and metastatic lesions and their relationship with immune infiltration. Finally, the deconvolution of these spatial niches in a large cohort of bulk RNA samples revealed distinct clusters associated with patient survival.

## Material and Methods

### Data collection and metadata standardization

We integrated datasets that performed single-cell or single-nuclei RNA-seq on all cells without prior enrichment by cell sorting. Our data collection included samples from healthy donors, adjacent normal tissue, primary PDAC, and metastatic PDAC. For metadata standardization, we selected relevant features for reproducibility (such as batch, dataset, patient_id, sample_id, age, sex, stage, condition, treatment, drug, author annotation if available, and atlas annotation). We also specify if the samples were processed by scRNA or snRNA protocol.

### Data download

Pre-processed datasets were mostly acquired from GEO with the exception of Zhou’s dataset (downloaded from the HTAN repository) (20) and Peng’s dataset (obtained from NGDC access code CRA001160) (16). Steele (17), Lin (7), Lee (8), Chen (23), Hwang (19), Carpenter (22), Werba (21), and Elyada’s (18) datasets were acquired from GSE155698, GSE154778, GSE156405, GSE212966, GSE202051, GSE229413, GSE205013, and GSE129455 respectively. Bulk RNA from TCGA data and GSE253260 (31) were downloaded with the TCGAbiolinks package (32).

### Data processing, normalization, integration, and clustering

We followed recently established benchmarks for quality control. Removal of low-quality cells removal considering 3 main variables together (number of UMI, number of genes, and percentual of mitochondrial counts) (33). Cell filtering was applied within each dataset separately as follows: > 5 MAD (median absolute deviation) for both the number of UMI and the number of genes detected while > 3 MAD for the percentage of mitochondrial counts with a maximum cut-off of 30%. Potential doublets were identified and removed with SOLO (34) implementation from scVI-tools. We have implemented a standard procedure for data normalization. Single-cell analysis was performed with Scanpy (35). First, data was normalized with a global scale factor of 10.000 and log1p transformation. Then, we performed feature selection with the function sc.pp.highly_variable_genes (ngenes = 6000, flavor = “seurat_v3”, layer = “counts”, batch_key = “dataset”).

Datasets were integrated with scANVI (36). Since scANVI requires cell annotation and a pre-trained scVI model, we manually annotated Zhou_2022 and Werba_2023 separately and used them as *seed* datasets. Our scVI model was trained with the following parameters (n_layers = 2, n_latent = 30, batch = ‘sample_id’, gene_likelihood = ‘nb’) and a maximum of 500 epochs. For the scANVI model, we set the maximum number of epochs to 200. We extracted the batch-corrected embeddings from the scANVI model and ran both *sc.pp.neighbors* and *sc.tl.umap* with default parameters. Finally, we performed clustering with the Leiden algorithm (resolution = 1.5) and reannotated the clusters with established gene signatures from the literature.

### Copy-number variation score (CNV score)

Infercnvpy (https://github.com/icbi-lab/infercnvpy) is a plugin derived from the *inferCNV* library to infer copy number variation from single-cell transcriptomics data. We downloaded the GTF file from the Xena browser to obtain the chromosome location for each gene in our matrix. Then, we followed the *infercnvpy* tutorial and defined all non-epithelial cells as our reference.

### Differential expression analysis

We leveraged the versatility of the scVI (37) package and built a model to identify differentially expressed genes. We considered a gene to be differentially expressed only if it showed FDR < 0.1 and logFC > 0.25.

### Non-negative matrix factorization (NMF) for the identification of malignant gene programs

NMF was performed with the *GeneNMF* package (38). First, we split our Seurat object by batch (*e.g.* “sample_id”) and followed the package recommendations. We used the top 2000 highly variable genes. We ran multiple values of K (4–9) for each one of the objects. We set 50 as the maximum number of genes and 10 as the maximum number of metaprograms.

### Gene signatures with UCell

Gene signature scores were assessed with *UCell* (39), an R package, via the *AddModuleScore_UCell* function that calculates the score of the geneset for each cell individually.

### Gene-set enrichment analysis

We used all genes identified by each NMF program and differentially expressed genes identified by the Wilcoxon rank sum test and performed annotation with *clusterProfiler* (40) package. Our parameters were *ontology* = ’Biological Processes’ and *qvalueCutoff* = ‘0.05’.

### Gene-set variation analysis

We applied the gene-set variation analysis pipeline from the GSVA package (41). First, gene expression was normalized and log-CPM values were used as input following the recommendations. We ran *gsvaParam* with the parameters *maxDiff* set to TRUE and *kcdf* set to “Gaussian”.

### Identification of TME subtypes

For this analysis, we removed epithelial cells, endocrine cells, and cell types that were almost exclusive to a few datasets (neutrophils, mast cells, Schwann cells, plasma cells). We created a pseudobulk for each sample and applied GSVA to estimate the TLS score, signature obtained from Wang et al (42), and the 29 FGES from Bagaev et al (43). Next, we merged the TLS score with the relative abundance and performed median scale normalization. Then, we used this matrix and created an Anndata object. We performed PCA, with the function *sc.tl.pca* (svd_solver = ‘arpack’), followed by *sc.pp.neighbors* (n_neighbors = 20), *sc.tl.umap* (min_dist = 0.25, maxiter = 2500, spread = 2) and Leiden algorithm (resolution = 0.5).

### Cell-cell interactions with CellChat

We used our scRNA-seq data to predict potential cell-cell interactions with the *CellChat* (44) R package. For input, we provided our scRNA-seq matrix with all cell types. We used cell state annotation for B, CAF, CD4+T, CD8+T, dendritic cells, monocytes, and macrophage clusters. First, we constructed separate CellChat objects for each condition of interest as recommended in the tutorial. Afterward, we conducted a comparative analysis to infer differentially enriched ligand-receptor interactions within the TME. We visualized the outgoing signaling and incoming signaling strength of the cell types between the different conditions via the *netAnalysis_signalingRole_scatter* function. Furthermore, we identified the significantly enriched pathways L-R interactions across all conditions using the *netAnalysis_signalingRole_heatmap* function, setting the options *pattern =* “*all*” to aggregate the incoming and outcoming signaling strengths.

### Spatial deconvolution and spatial niches identification

RCTD (45) was used for spatial deconvolution. We used our single-cell atlas (only PDAC and MET samples were kept) as our reference dataset to perform deconvolution. The spatial transcriptomics datasets used were Zhou’s dataset (including 15 samples of primary PDAC untreated or treated) and Khaliq’s dataset (46) (including 3 normal pancreata, 10 primary PDAC, 12 liver metastasis, and 5 metastatic lymph nodes). Doublet_mode was set to “full”. Finally, we extracted the normalized values from the RCTD object and built an Anndata object. After verifying the elbow plot, we applied k-means (k = 20) and recovered the spatial niches.

### Deconvolution for cell type prediction and recovery of spatial niches in bulk RNA samples

We performed bulk deconvolution with BayesPrism (47). Initially, we conducted the deconvolution of our 52 cell types described in the atlas in a large cohort of bulk samples. We created the prism object with our single-cell reference matrix containing only the signature genes identified for each population. *new.prism* function was used as follows: *reference* = “sc_matrix”, *mixture* = “bulk_matrix”, *input.type* = “count.matrix”, *key* = “Malignant ductal cells”, *outlier.cut* = 0.01, *outlier.fraction* = 0.1. For spatial niche recovery in bulk samples, first, we performed feature selection by retaining only genes with logFC > 0.75 | p.value < 0.05 | score > 10 from the Wilcoxon rank-sum test (*sc.tl.rank_genes_groups*). Then, we used DeconvoBuddies (48) to find niche marker genes with the *get_mean_ratio* function and retained genes with a mean ratio above 0.75. Finally, we combined those two approaches and retained only the shared genes. We extracted the SCT corrected matrix containing only the marker genes and created a prism object with only a single change in the parameters above (*key* = NULL).

### Bulk RNA groups

We ran GSVA with the 29 FGES from Bagaev on bulk samples. After that, we merged this matrix of GSVA scores with the spatial niche proportion matrix and performed Z-score normalization. Then, we applied hierarchical clustering to the Pearson correlation among patients based on the spatial niche proportion and GSVA scores (distance = “Euclidean”, method = “ward.D2”). To determine the ideal number of clusters, we used an automated approach from the dynamicTreeCut R package (function *cutreeDynamic*) (49).

### Survival analysis

The Kaplan–Meier survival curves were plotted using the R packages “survival” and “survminer”. The Coxph test was used to determine the hazard ratio (HR) and 95% confidence interval.

### Statistical analysis

For comparing continuous variables, we used the two-sided Wilcoxon rank-sum test or the Kruskal-Wallis test. Multiple testing corrections were performed where necessary using the Benjamini-Hochberg method.

## Results

### Building a robust single-cell atlas for PDAC

In the present study, we integrated 10 publicly available datasets into a comprehensive PDAC single-cell atlas containing 259 samples from 202 patients, totalizing 941.608 high-quality cells after rigorous quality control. Our atlas includes data from 22 samples of normal pancreatic tissue, 8 samples of adjacent normal tissue, 211 samples of primary PDAC, and 18 samples of metastatic PDAC. We next leveraged our refined cellular annotation to use as a reference for spatial deconvolution of two large datasets of spatial transcriptomics to comprehend the spatial distribution and co-localization of 52 fine-grained cell types. Finally, we performed deconvolution of a large cohort of bulk samples to assess the clinical impact of the spatial niches identified in our atlas (**Figure 1A**).

**Figure 1.**
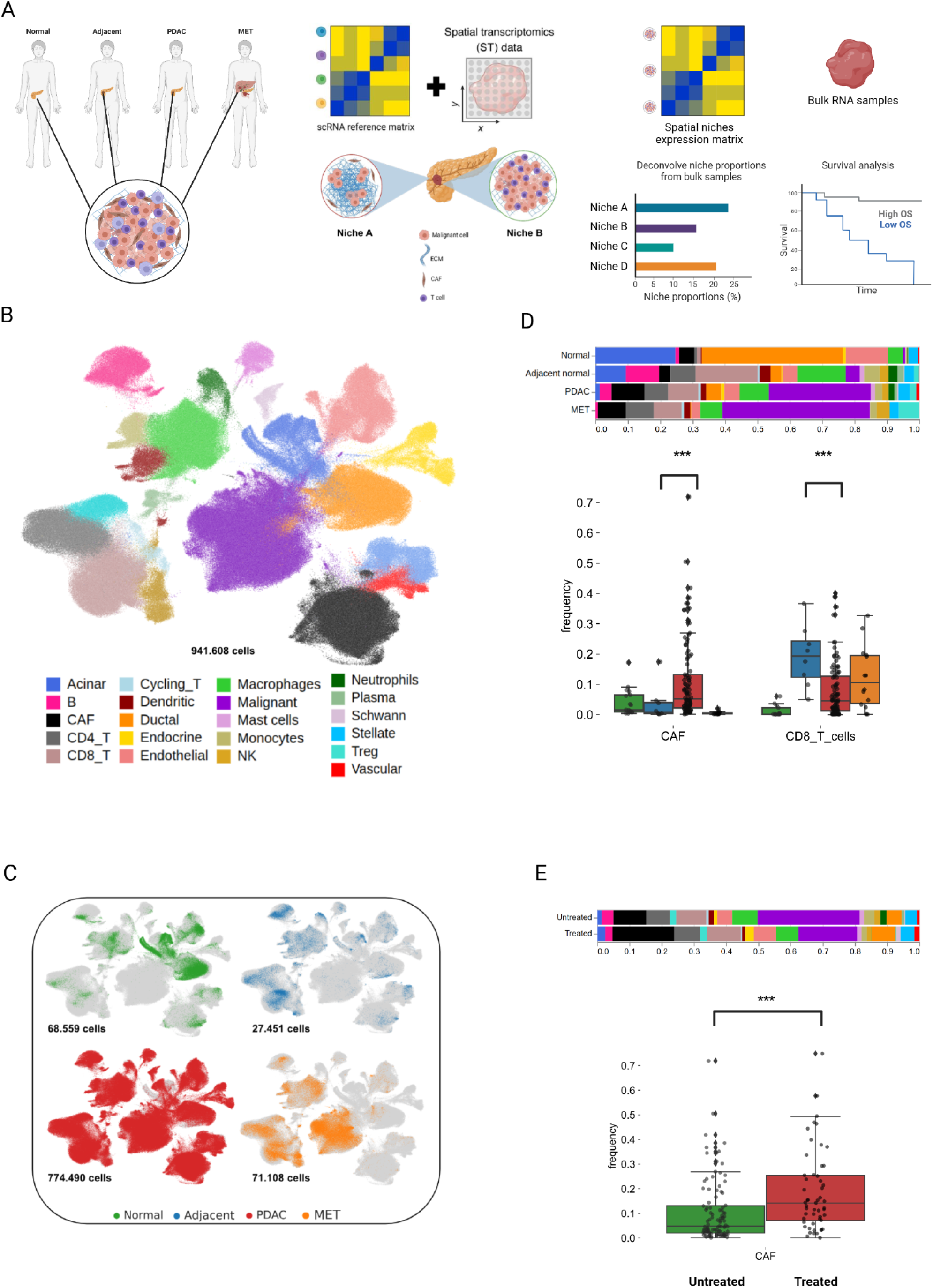
PDAC atlas overview -. (**A**) Overview of the study workflow; (**B**) UMAP projection of the atlas; (**C**) UMAP projection colored by each biological condition present in the atlas; (**D**) Top - Stacked bar chart for Normal, Adjacent normal, PDAC and MET cell type composition. Bottom - Wilcoxon test for CAF and CD8+T proportions. (p < 0.001; Wilcoxon two-sided test); (**E**) Top - Stacked bar chart for untreated and treated PDAC cell type composition. Bottom - Wilcoxon test for CAF proportions. (p < 0.001).

One of the main challenges faced when integrating multiple datasets is the batch effect. Most methods can easily handle batch effect, however they may not preserve biological heterogeneity (50). Hence, we implemented scANVI which is one of the top performers for large-scale data integration and preserves biological heterogeneity (detailed in Methods). Nearly 1 million cells were annotated in 21 coarse cell types based on a rigorous revision of established canonical signatures in the literature (**Figure 1B** and **Supplementary Figure 1A**). For malignant cell annotation confirmation, we applied *inferCNVpy* to identify cells with CNV alterations (**Supplementary Figure 2B**). As expected the malignant cells cluster presented a higher inferCNV score than acinar and normal ductal cells (**Supplementary Figure 2C**). Besides, ductal cells derived from normal pancreas samples formed a distinct cluster and did not overlap with malignant cells (**Figure 1C**). In an alternative approach to previous works (51,52), we decided to integrate scRNA and snRNA-seq data to generate a comprehensive reference atlas considering these modalities can introduce a bias toward capturing one particular cell type with higher efficacy than the other (*e.g* immune cells are well represented in scRNA while complex cell types benefit the most from snRNA). Our pipeline was able to mitigate any significant batch effect between datasets generated with scRNA or snRNA (**Supplementary Figure 1B**). In agreement with the literature (53–56), we observed a higher recovery of stromal and epithelial cells in snRNA whereas scRNA exhibited a higher fraction of T cells (**Supplementary Figure 2A**).

Next, we explored the differences in cellular composition across distinct microenvironments in the pancreas. As expected, normal pancreatic tissue samples were enriched with acinar and normal ductal cells, with minimal presence of immune cells. In contrast, adjacent tissue showed pronounced immune infiltration, particularly T lymphocytes (CD4+ and CD8+) and B lymphocytes, indicating immunological activity in this microenvironment. Primary tumors were marked by enrichment of CAFs, typical of the desmoplastic reaction in PDAC. Additionally, primary tumors had fewer CD8+ T cells than adjacent tissue. Metastatic lesions were characterized by a microenvironment with scarce stromal cells and rather dominated by malignant cells with a modest immune cell infiltration (**Figure 1D** and Supplementary Figure 1C). Considering that our atlas does not include paired samples from the different microenvironments, we used a dataset (57) with paired adjacent tissue and primary tumor samples to validate this finding. After unsupervised clustering and cell annotation (Supplementary Figure 2D), we compared the composition of adjacent tissue and PDAC between paired samples. We found a higher fraction of CD8+ T cells in adjacent tissue compared to primary tumors confirming our results from the atlas (Supplementary Figure 2E). Finally, we evaluated the dynamics of cellular composition between treated and untreated PDAC samples. As expected, treated samples had fewer malignant cells. We also observed that treated samples exhibited a significant increase in the proportion of CAFs consistent with prior studies (9,19,58) (**Figure 1E**). Altogether these results highlight the poor immune infiltration in PDAC and TME remodeling after chemotherapy.

### CD8+ T subset signatures differ between treated and untreated PDAC microenvironments

Because we found a significant enrichment of CD8+ T cells in the adjacent tissue, we aimed to refine our characterization of cytotoxic lymphocytes, reclustering our subset of CD8+ T cells. Based on canonical marker investigation, differentially expressed genes, and public signatures, we identified eight cellular states of CD8+ T cells: NL (naive-like), CM (central memory), MAIT (mucosal-associated invariant T cell), EM (effector memory), TRM (tissue-resident memory), TEMRA (terminally differentiated effector memory cells), TPEX (progenitor exhausted cells), and TEX (exhausted cells) (**Figure 2A**).

**Figure 2.**
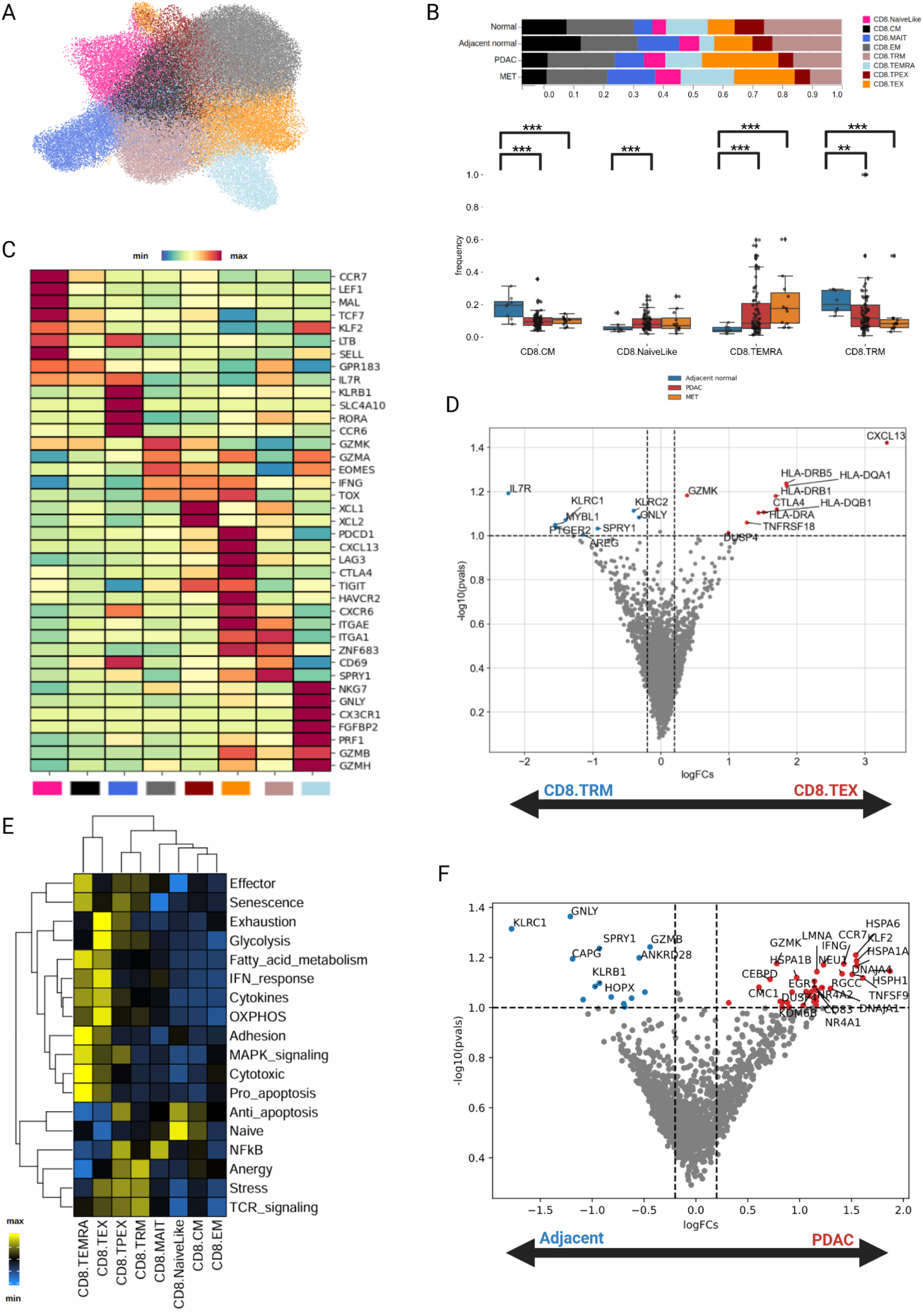
CD8-TRM cells are enriched in adjacent tissue. (**A**) - UMAP projection of 8 clusters of CD8+T cells; (**B**) Top - Stacked bar chart for Normal, Adjacent normal, PDAC, and MET cell type composition. Bottom - Wilcoxon two-sided test for CD8+T clusters. (**\*\*\*** = p < 0.001; **\*\*** = p < 0.01); (**C**) - Clustered averaged expression of markers used for annotation (Z-score normalized expression); (**D**) - Volcano plot of differentially expressed genes between CD8.TRM (left) and CD8.TEX (right) (FDR < 0.1); (**E**) - Heatmap exhibiting expression of CD8+T functional signatures (Z-score normalized values); Volcano plot of differentially expressed genes between adjacent CD8+T cells (left) and PDAC CD8+T cells (right) (FDR < 0.1).

The CD8-NL cluster was identified by the high expression of CCR7, SELL, and TCF7 whereas the CD8-CM cluster exhibited decreased expression of these markers and higher expression of the TCR signature, indicating that they had already undergone activation. CD8.MAIT cluster showed high expression of KLRB1, SLC4A10, RORA, and CCR6 consistent with previous annotations. CD8-EM cells demonstrated high expression of GZMK, GZMA, and EOMES. CD8.TRM cells were identified by the high expression of canonical tissue retention markers, such as CD69, CD49a (ITGA1), HOBIT (ZNF683), SPRY1, and CD103 (ITGAE). CD8-TEX showed high expression of some tissue retention markers, along with expression of PDCD1, CTLA4, CXCL13, and other immune checkpoint proteins (**Figure 2C**). However, CD8-TEX also exhibited a high IFN signature, indicating that they were not fully dysfunctional (**Figure 2E**). Considering the overlap of some features between CD8-TEX and CD8-TRM clusters we asked which features distinguish these two clusters. Differential expression analysis revealed high expression of CXCL13, CTLA4, DUSP4, GZMK, and MHC-II genes in the CD8-TEX cluster, while the CD8-TRM cluster showed high expression for IL7R, AREG, GNLY, KLRC1, and KLRC2 (**Figure 2D**). CD8-TPEX cells expressed the transcription factor TOX, IFNG, and chemokines XCL1 and XCL2 responsible for dendritic cell recruitment. TEMRA cells displayed high expression of cytotoxicity-related genes such as GNLY, NKG7, PRF1, and GZMB (**Figure 2C**).

We next sought to determine if any specific CD8+T subset was enriched in the adjacent microenvironment. CD8-TRM cluster was the predominant population in adjacent tissue and was present in higher proportions compared to the primary tumor, which, in contrast, had a higher fraction of CD8-NL cluster. To confirm this result in paired samples, we reclustered CD8+T cells and annotated TRM cells based on the expression of CXCR6, ITGAE, ITGA1, and ZNF683 (Supplementary Figure 3D-E). We confirmed a higher composition of CD8-TRM in the adjacent tissue of matched samples (Supplementary Figure 3F and H). CD8-CM cluster was also enriched in adjacent tissue compared to malignant tissue, while CD8-TEMRA was enriched in the malignant tissue (**Figure 2B**). Differential expression analysis of CD8+T cells between matched samples revealed higher expression of cytotoxic genes, such as GZMB, GNLY, KLRC1, and KLRC2, and lower expression of heat-shock genes, IFNG, CCR7, and KLF2 in adjacent tissue (**Figure 2F**). Furthermore, we compared the expression of cytotoxic and effector genes in the CD8-TRM cluster from matched samples. We found increased expression of PDCD1, GZMB, PRF1, GNLY, CD27, and CXCR3 in TRM cells from adjacent tissue compared to the primary tumor (Supplementary Figure 3G). These findings indicated a potential higher CD8+T cell response active in the adjacent, but not in the malignant tissue.

For many tumors, chemotherapy has a strong immunogenic effect (59–61) however this has not been fully elucidated in PDAC. We thus evaluated if any impact of chemotherapy could be detected in our samples. We identified that, while an enrichment of CD8-TEX and CD8-TEMRA clusters exists in untreated PDAC patients, treated PDAC patients showed a higher proportion of CD8-TPEX and CD8-TRM clusters (Supplementary Figure 3A-B). Because CD8-TEX cells are characterized by high expression of interferon signaling and cytotoxic activity (**Figure 2E**) we analyzed the differential expression between CD8-TEX clusters from treated and untreated patients. Higher expression of CXCL13, CTLA4, and ENTPD1 was detected in untreated patients - a signature currently considered a predictor of immunotherapy response in pan-cancer analyses (62,64). In contrast, treated patients showed high levels of GZMK, EOMES, and KLRG1 (Supplementary Figure 3C), which has been previously associated with a poor immune response in melanoma (64). In summary, treatment in PDAC leads to a less functional CD8+T cell response in malignant tissue.

### iCAF proportion is associated with a good prognosis in PDAC

The desmoplastic reaction is one of the main hallmarks of pancreatic cancer and plays a major role in tumor initiation, tumor progression, therapy response, and prognosis (65–67). Here, we isolated stromal cells and reclustered the data to obtain fine-grained populations.

We identified four clusters of CAF (DP_CAF, iCAF, myCAF, and Cycling_myCAF), two clusters of stellate cells (MYH11+ and RGS5+), five clusters of endothelial cells (capillary, lymphatic, ACKR1+, CXCL12+ and RGS5+), one cluster of Schwann cells and one cluster of vascular smooth muscle cells (**Figure 3A**). Cycling_myCAF population was marked by expression of MKI67, FN1, STMN1; DP_CAF (double positive) showed expression of both iCAF and myCAF together with overexpression of CXCL14, SFRP1, SFRP4. This population was identified in a prior study (68). iCAF was identified by high levels of C3, C7 and PTGDS whereas myCAF overexpressed CTHRC1, COL11A1 and MMP11 (**Figure 3C**). Bulk deconvolution with BayesPrism (47) unraveled increased survival for patients with higher fractions of iCAF, however, there was no correlation between myCAF and survival (**Figure 3C-D**). These results are consistent with a recent study that observed increased survival in patients with a higher fraction of iCAF (69).

**Figure 3.**
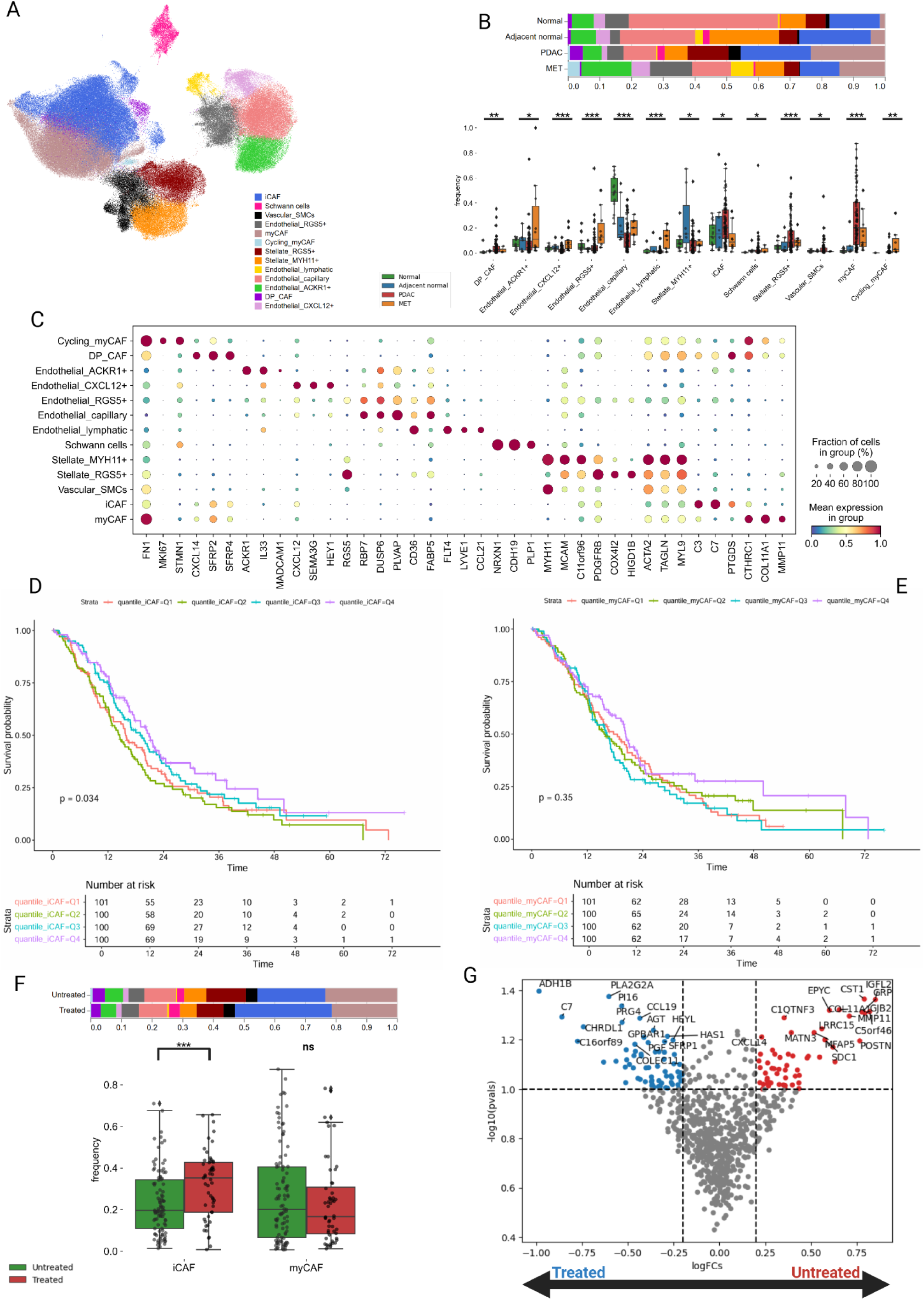
iCAFs correlate with improved survival in PDAC. (**A**) - UMAP projection of the stromal cells; (**B**) - Top - Stacked bar chart for Normal, Adjacent normal, PDAC, and MET cell type composition. Bottom - Wilcoxon two-sided test for stromal clusters. (**\*** = p < 0.05; **\*\*** = p < 0.01,**\*\*\*** = p < 0.001 ); (**C**) - Dot plot with marker genes for each cluster; (**D**) - Kaplan–Meier plot showing the associations between the iCAF cluster with overall survival in the bulk cohort (stratified by quantile). P values were calculated using the log-rank test; (**E**) - Kaplan–Meier plot showing the associations between the myCAF cluster with overall survival in the bulk cohort (stratified by quantile). P values were calculated using the log-rank test; (**F**) - Top - Stacked bar chart for untreated (green) and treated (red) PDAC CAFs composition. Bottom - Wilcoxon test for CAF clusters. (**\*** = p < 0.05; ns = non significant); (**G**) - Volcano plot of differentially expressed genes between untreated CAF (right) and treated CAF (left) (FDR < 0.1).

Stellate MYH11+ had high expression of MYH11, MCAM, and C11orf96 and shared other markers with vascular smooth muscle cells. This population was previously observed around vessels (70). Whereas Stellate RGS5+ cells were marked by high levels of RGS5 and hypoxia-related genes such as COX4I2, and HIGD1B. Schwann cells were also present in the atlas and were identified based on expression of NRXN1, CDH19, and PLP1; Vascular smooth muscle cells displayed high levels of MYH11, ACTA2, and MYL9 (genes involved in actin or myosin pathways) (**Figure 3C and Supplementary** Figure 4).

Endothelial capillary (or endothelial stalk cells) represented the majority of the population of endothelial cells and showed high expression of PLVAP, CD36, and FABP5. On the other hand, lymphatic endothelial cells were marked by LYVE1, FLT4, and CCL21 (genes related to immune cell chemoattraction). The endothelial ACKR1+ population was marked by high levels of ACKR1, IL33, and MADCAM1 - known markers for endothelial cell activation and recruitment of immune cells (71). Endothelial CXCL12+ showed high expression of CXCL12, SEMA3G, and HEY1 indicating a potential role in neovascularization and leukocyte transmigration (72). Finally, we also identified a cluster of endothelial cells (Endothelial RGS5+) that is likely undergoing endoMT (endothelial mesenchymal transition) as evidenced by expression of PDGFRB and ameboidal type cell migration pathway (**Figure 3C and Supplementary** Figure 4).

Then, we explored cell type composition differences between the conditions. The normal pancreas was highly enriched for endothelial cells, mainly endothelial capillary, and lower fraction of CAFs compared to other conditions as expected. Endothelial capillary cells were the predominant population in the normal pancreas and it was already severely decreased in the adjacent tissue as a result of tissue cancerization. As expected, primary PDAC tumors showed an increase in CAFs especially myCAF responsible for desmoplastic reaction. Moreover, we also observed a higher fraction of Stellate RGS5+ indicating that these cells are being activated. Metastatic lesions had significantly fewer stromal cells compared to primary tumors in general, although we observed a decrease in endothelial capillary population and an increase in Endothelial_ACKR1+ and RGS5+ (**Figure 3B**). We previously observed enrichment for CAFs in treated patients compared to untreated ones, hence we asked whether there were significant changes in CAFs composition. Treated patients exhibited a higher proportion of iCAF which was confirmed by differential expression analysis (**Figure 3F**). Additionally, we also noticed a high expression for CCL19 - chemokine responsible for T-cell attraction - and low expression of LRRC15 - that was linked to immunotherapy resistance (73,74) (**Figure 3G**). Collectively, these findings reveal a shift in CAFs composition and transcriptional profile in treated PDAC.

### Malignant cells MHC-II+ negatively correlate with T-cell infiltration and are enriched in metastatic lesions and treated PDAC

Malignant cells in PDAC present high transcriptional heterogeneity and previous studies with bulk RNA identified two main subtypes: classical and basal (75–77). These subtypes were recovered with scRNA-seq (78–80) and Raghavan et al discovered a hybrid population sharing both transcriptional programs (10). Recently, Hwang et al identified three new subtypes in addition to previous classifications (19). Here, we performed NMF analysis revealing eight transcriptional programs (from here on called MP) (**Figure 4A**).

**Figure 4.**
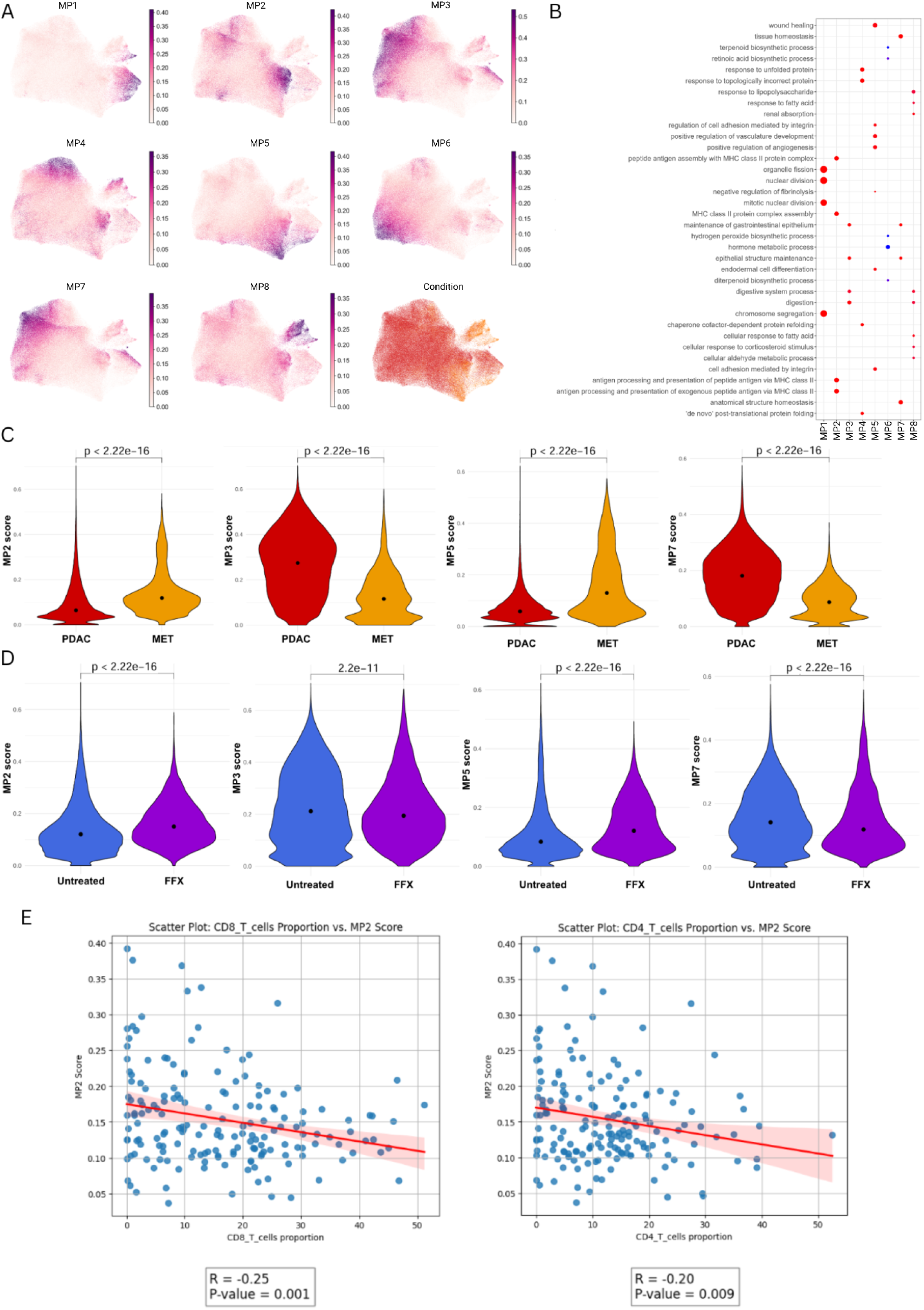
Malignant cells MHC-II+ negatively correlate with T-cell infiltration in primary and metastatic PDAC. (**A**) - UMAP feature plot for MP (Metaprograms) scores; (**B**) - Gene ontology for malignant MP’s. Rows indicate the pathways obtained from the BP (Biological Process) of GO. Columns represent the malignant MP’s; (**C**) - Violin plot evaluating for MP2, MP3, MP5, and MP7 among PDAC and MET. Dot indicates the median expression; (**D**) - Violin plot evaluating for MP2, MP3, MP5, and MP7 among untreated PDAC and treated PDAC. Dot indicates the median expression; (**E**) - Left - Scatter plot showing the correlation between MP2 score and CD8+T proportion; Right - Scatter plot showing the correlation between MP2 score and CD4+T proportion.

MP1 was predominantly represented by cell cycle genes (**Figure 4B**). MP2 was enriched for MHC-II genes and it was not previously described. MP2 score negatively correlated with CD4 and CD8+T cell infiltration (**Figure 4E**). MP3 showed high expression of classical markers and overlapped with classical signatures previously identified by Moffitt and others (75–77). MP7 also exhibited overlap with classical signatures (Supplementary Figure 5A). MP4 was associated with stress response signature based on high expression of heat shock proteins. MP5 exhibited upregulation of FN1, SNAI2, CAV1, and VIM overlapping with basal signatures (75–77) and resembling EMT-state cells (Supplementary Figure 5B-C). MP6 showed the expression of genes related to metabolism pathways. MP8 was enriched in response to LPS (Lipopolysaccharide) and response to fatty acid pathways, in addition, it was prevalent in metastasis (**Figure 4A-B**). We also observed a positive correlation between MP2 and MP5 scores suggesting that downregulation of MHC and EMT are connected (Supplementary Figure 6A).

Then, we compared the MP scores between malignant conditions and noticed higher scores for MP3 and MP7 (classical phenotype) in primary tumors, whereas metastatic lesions exhibited higher scores for MP2 and MP5 (**Figure 4C**). These signatures were also increased in treated primary tumors compared to untreated uncovering EMT and MHC-II upregulation as potential mechanisms of therapy resistance and immune evasion (**Figure 4D**). These findings are consistent with previous reports that observed a poor chemotherapy response and reduced survival for the basal subtype (9,75–77,80,81).

Next, we focused on patients with matched snRNA and spatial transcriptomics (10x Visium) from Zhou’s dataset and explored the spatial distribution of these eight phenotypes. To annotate the spots, we considered only spots with a fraction of malignant cells above 0.7 and then assigned them based on the highest MP score. We found that MP5 was preferentially located at the tumor border compared to MP3 and MP2, consistent with their basal/EMT-like phenotype (Supplementary Figure 7). MP3 mapped mostly to tumor glands confirming their classical phenotype. MP2 presented a scattered spatial distribution unlike MP3 and MP5, except for patient HT264P1 which the majority of the spots were annotated as MP2 (Supplementary Figure 7). These findings correlate with the single-cell results as patients with high expression of particular MP also had most of the malignant spots assigned to respective MP (*e.g.* HT264P1 enriched for MP2, HT360P1 enriched for MP5, HT231P1, and HT259P1 enriched for MP3) (Supplementary Figure 6C **and Supplementary** Figure 7). We could not assign spots for patient HT288P1 because the slice had a low tumor fraction. Finally, we obtained another spatial dataset (46) that contains primary PDAC, metastatic lymph nodes, and liver metastasis samples to validate our findings. Applying the workflow described earlier to retain only spots enriched in malignant cells, we identified a higher MP2 score in metastatic lymph nodes and liver metastasis compared to primary PDAC confirming our previous results (Supplementary Figure 6B). Overall, our data uncovered a new transcriptional subtype of malignant cells negatively correlated with immune infiltration and is consistent with the literature showing that the basal signature is enriched after treatment (9,10,58).

### Single-cell composition of the TME reveals distinct PDAC tumor immune phenotypes

To comprehend the bigger picture of TME heterogeneity in PDAC, we stratified the patients based on cell type composition and a TLS score generated with GSVA (more details in the methods section). For the TLS score, we first converted our single-cell matrix to pseudobulk by patient. After that, we ran GSVA for a 30-gene signature for TLS (42) and recovered the score for each patient.

We performed dimensionality reduction followed by unsupervised clustering revealing six distinct phenotypes: Normal-like (low presence of CAF, high proportion of endothelial and stellate cells, together with acinar and non-malignant ductal cells); Desertic (poor immune infiltration and predominance of malignant cells followed by CAF enrichment); Immune-Fibrotic (high proportion of CAF and CD4-CD8 T cells); Lymphoid-Rich (enriched for adaptative immune response including B cells, CD4+T, CD8+T and TLS signature); Immune-Mixed (rich for CD4 T and monocytes) and Myeloid-Rich (enriched for dendritic cells and mainly macrophages) (**Figure 5A-B and Supplementary** Figure 9A).

**Figure 5.**
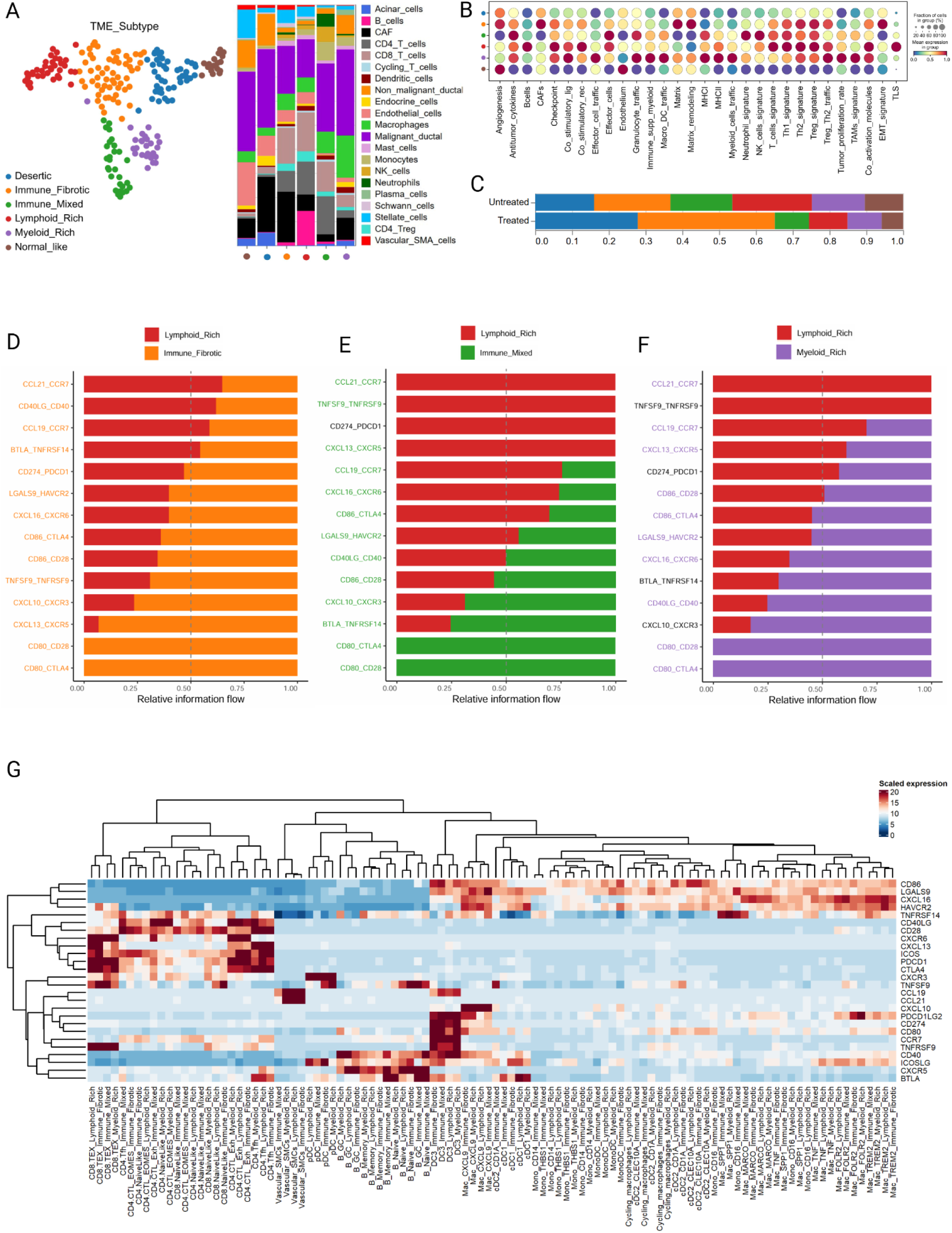
Tumor immune phenotypes in PDAC. (**A**) - Left - UMAP projection of the six TME subtypes; Right - Stacked bar chart showing the cell type composition across the TME subtypes; (**B**) - Dot plot exhibiting the 29 FGES from Bagaev; (**C**) - Bar chart of the TME subtype composition among treated and untreated PDAC patients; (**D-F**) - Bar plot for ligand-receptor pair showing the relative flow information; (**G**) Heatmap showing the expression for each ligand and receptor across Lymphoid-Rich, Immune-Fibrotic, Immune-Mixed and Myeloid-Rich.

We applied the FGES signature from Bagaev (43) and identified increased expression of immune checkpoint genes, co-stimulatory receptor, co-activation molecules, and Th1 signatures in Lymphoid-Rich patients (**Figure 5B**). As expected, Immune-Fibrotic patients showed higher expression of matrix, matrix remodeling, EMT, and CAF signatures. Strikingly, the Lymphoid-Rich group showed lower expression of MHC-I and MHC-II signatures than Myeloid-Rich. The latter group also displayed higher expression of effector cell traffic, co-stimulatory ligand, macrophages-dendritic cells traffic, and immune suppressive myeloid population showing both anti-tumoral and pro-tumoral features (**Figure 5B**). The majority of treated patients were classified as either Desertic or Immune-Fibrotic highlighting the impact of chemotherapy in remodeling the TME architecture towards a fibrotic environment with or without immune infiltration (**Figure 5C**). B cells from the Lymphoid-Rich microenvironment exhibited upregulation of APC and coactivation signature suggesting a potential role in antigen presentation (Supplementary Figure 8A-D).

Next, we focused on the differences in cell-cell interactions between the TME subtypes with immune infiltration. Among the four subtypes (Lymphoid-Rich, Immune-Fibrotic, Immune-Mixed, and Myeloid-Rich) we found that the CD8-TEX cluster was the main target of incoming signaling in Lymphoid-Rich and Immune-Fibrotic, but not in Immune-mixed and Myeloid-Rich whose main target was CD8-TRM cluster (Supplementary Figure 9B). For all subtypes, myCAF and iCAF dominated the outgoing signaling highlighting their strong influence on extracellular communication (Supplementary Figure 9B). Then, we explored the relative signaling flow of immune ligand-receptor pairs and found enrichment for CCL19-CCR7 and CCL21-CCR7, interactions involved with homing and recruitment of dendritic and T-cells and important for the development of TLS (82–85), in Lymphoid-Rich compared to the other subtypes justifying the higher infiltration of T-cells in this group (**Figure 5D-F**). These pairs were also higher in Immune-Fibrotic compared to Immune-Mixed and Myeloid-Rich (Supplementary Figure 9C-D). Surprisingly, CD80-CD28 and CD86-CD28 pairs were less pronounced in the Lymphoid-Rich subtype. The CXCL13-CXCR5 pair, responsible for B cell attraction and formation of TLS (86–88), showed a strong signal in Lymphoid-Rich, however, this pair exhibited a stronger signal in the Immune-Fibrotic subtype (**Figure 5D-G**). Strikingly, several ligand-receptor pairs were enriched in Immune-Fibrotic compared to Immune-Mixed and Myeloid-Rich (*e.g.* CD80-CD28, CD86-CD28, and CXCL13-CXCR5) suggesting a stronger antitumor immune response despite a fibrotic environment (**Figure 5G and Supplementary** Figure 9C-D).

### Spatial deconvolution uncovers conserved cellular niches across PDAC patients

Finally, to gain insights into the spatial architecture and cellular communities we integrated 2 datasets (20,46) of spatial transcriptomics (10xVisium). We applied RCTD (45) to infer cell-type proportions per spot using our single-cell atlas as a reference (Supplementary Figure 10A-G). Next, we created an Anndata object with our spot-level cell-type composition matrix and performed standard procedures (optimal number of clusters determination, k-means, and UMAP projection). We obtained 19 clusters (from here on called niches) based on cell type composition across 4 different biological conditions (normal pancreas, primary PDAC, metastatic lymph nodes, and liver metastasis) (**Figure 6A-D**).

**Figure 6.**
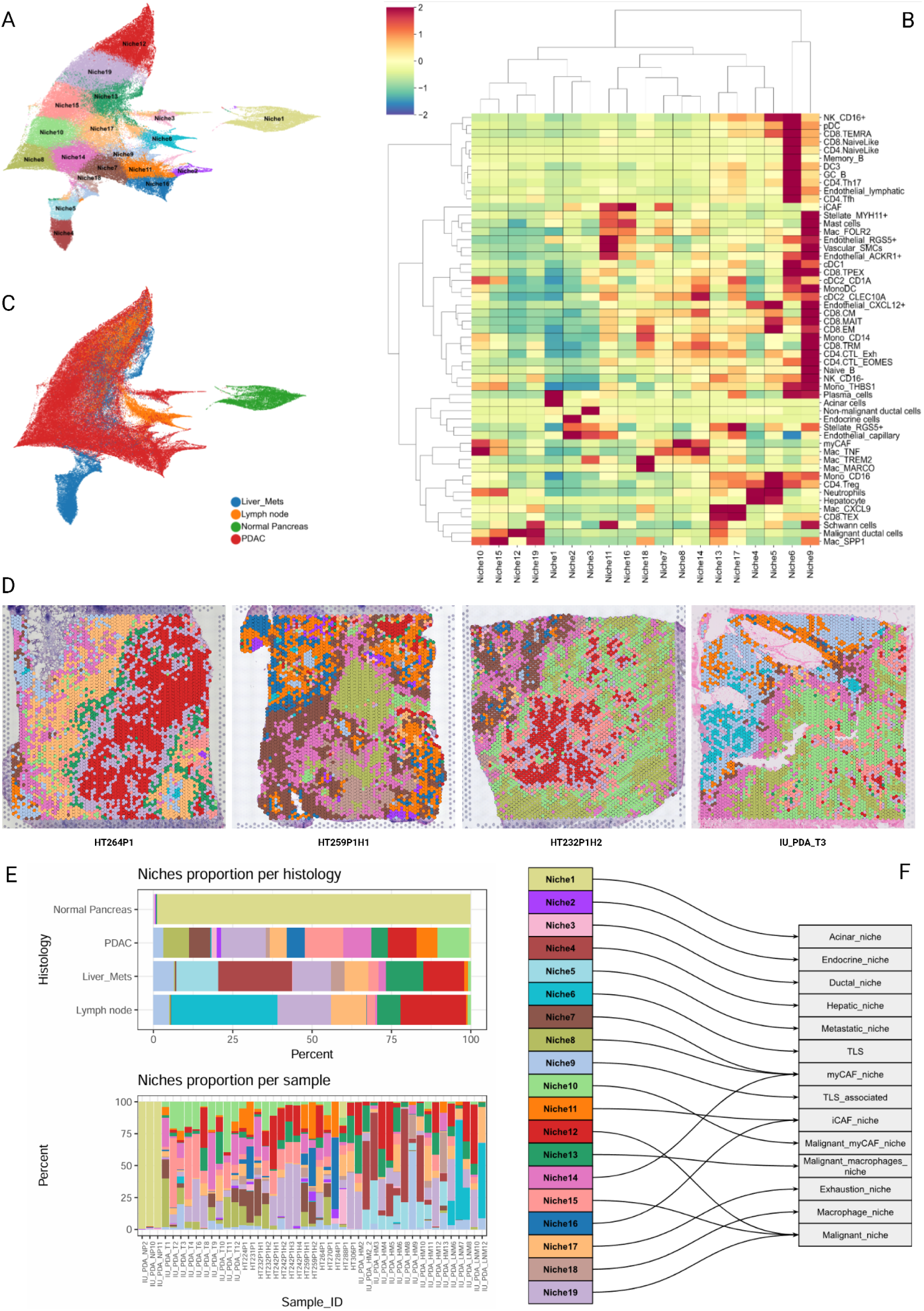
Spatial deconvolution uncovers conserved cellular niches across PDAC patients. (**A**) - UMAP projection colored by the 19 spatial niches; (**B**) - Heatmap showing mean cell type composition per spatial niche (Z-score normalized); (**C**) - UMAP projection colored by biological condition; (**D**) - Spatial distribution of the 19 niches in 4 representative PDAC samples; (**E**) - Top - Stacked bar chart showing niche composition per biological condition; Bottom - Stacked bar chart showing niche composition per sample; (**F**) - Niche annotation.

Niche1 was enriched for acinar cells and was predominant in normal pancreas slides and adjacent normal tissue. Niche2 and Niche3 were dominated by endocrine and normal ductal cells respectively (**Figure 6B and Supplementary** Figure 11B). Niche4 and Niche5 were exclusive to liver metastasis; the first was mostly composed of hepatocytes resembling normal anatomical areas whereas the latter showed enrichment for malignant ductal cells and an increase in inflammation evidenced by a higher fraction of different phenotypes of macrophages, CAFs, and stellate cells (**Figure 6B and Supplementary** Figure 11B).

Niche6 was marked by immune cells (germinal center B cells, follicular helper T cells, cDC1, DC3, memory B cells, and endothelial lymphatic cells). This niche was mostly identified in lymph node sections, but it was also present in a few tumors that contained tertiary lymphoid structures (TLS) (**Figure 6B-E**). We recovered a TLS-associated region, Niche9, enriched for plasma cells, effector populations of CD4 and CD8 T cells, and stromal-associated cells (*e.g.* stellate cells, and endothelial cells) along monocytes and macrophages (**Figure 6B and Supplementary** Figure 11B).

Niches 7, 11, and 16 were associated with fibrotic areas and were mainly composed of iCAF. We also observed an enrichment for Schwann cells in Niche11 suggesting a potential interaction between these populations (89). Niches 8 and 14 mapped to dense collagen areas and as expected were dominated by the presence of myCAF, although in the latter we observed a small infiltration of CD8-TRM and macrophages. Niche10 was linked to tumor stroma interface zones and it was rich in malignant cells and myCAF. This agrees with other studies that showed a proximity between myCAF and malignant cells (90).

Niches 12, 15, and 19 represented spots associated with tumor areas and composed mostly of malignant cells with the latter enriched for Schwann cells possibly related to the perineural invasion (91) (**Figure 6B and Supplementary** Figure 11B). Niches 13 and 17 revealed a co-occurrence of malignant cells with immune cells; in Niche13, macrophages (particularly Mac_CXCL9) were associated with malignant cells, whereas in Niche17 we observed an overlap of CD8-TEX with Mac_CXCL9 accompanied by a decrease in malignant cells proportion likely to be antitumoral activity based on high expression of GZMB, PRF1, and CD27 (**Figure 6B and Supplementary** Figure 11A-D). Finally, Niche18 was marked by distinct populations of immunosuppressive macrophages such as Mac_TREM2 and Mac_MARCO (**Figure 6B**). Considering a certain degree of cell type composition similarity between the niches, we decided to group Niches 12, 15, and 19 into “Malignant_niche”, Niches 7, 8, and 14 into “myCAF_niche”, and Niches 11 and 16 into “iCAF_niche” (**Figure 6F**).

After that, we combined two methods to obtain unique signatures for each niche (see Methods for details). Our next goal was to investigate if these spatial niches could have a clinical impact. We applied deconvolution with BayesPrism (47) to 2 cohorts of bulk RNA-seq. We also performed GSVA (41) with the 29 FGES from Bagaev (43) to obtain a refined molecular characterization of the TME. Then, we merged the matrices from deconvolution (niche composition per sample and GSVA scores) and performed Z-score normalization followed by hierarchical clustering revealing 9 clusters (**Figure 7A**).

**Figure 7.**
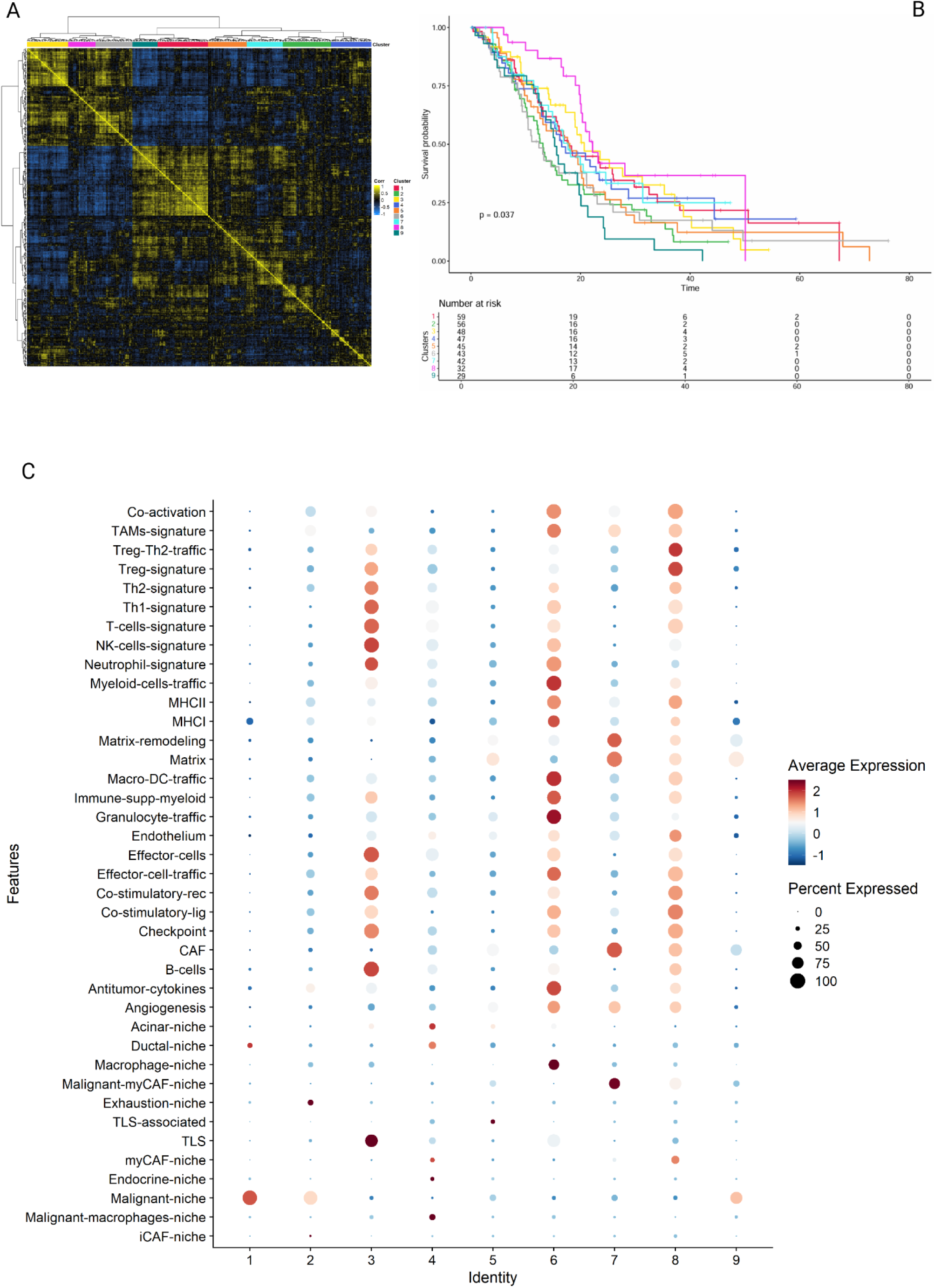
Niche composition and TME features predict clinical outcome. (**A**) - Heatmap for Pearson correlation across 401 bulk samples revealing 9 clusters; (**B**) - Kaplan-Meier survival curve among the 9 bulk clusters; (**C**) - Dot plot showing all 39 features scores per cluster.

Cluster 1 exhibited a high score for the malignant niche and downregulation of T-cell and Effector cell signatures, resembling a desertic phenotype. Cluster 2 presented a high score for the malignant niche, just like Cluster 1, but it also showed enrichment for TAM and Treg signatures (**Figure 7B-C**). Cluster 3 was identified by high TLS and effector cell scores yet, this antitumor response was damped by upregulation of immunosuppressive features such as high neutrophils and Th2 signatures and downregulation of MHC pathways (**Figure 7C**). Cluster 4 displayed a high score for the malignant-macrophage niche, upregulation of CAF signature, and downregulation of MHC-I and co-activation signatures. Cluster 5 exhibited enrichment for CAF, Matrix, Matrix-remodeling, Angiogenesis, and low expression of immune effector signatures (**Figure 7C**). Cluster 6 was enriched for malignant and macrophage niches accompanied by high scores of immunoregulatory signatures hindering successful antitumor response. Cluster 7 showed enrichment for malignant, malignant myCAF niches and downregulation of T cell, effector cell, B cell, and Th1 signatures (**Figure 7C**). Cluster 8 was enriched for MHC-II, immune checkpoint, and coactivation scores, although exhibited upregulation for Treg signature and myCAF niche, patients from this cluster had high overall survival (**Figure 7B-C**). Cluster 9 was highly enriched for the malignant niche and presented significant downregulation of immune pathways contributing to a poor prognosis (**Figure 7B-C**).

## Discussion

There are multiple single-cell studies in PDAC, therefore the urge to integrate all these datasets into a robust single-cell and spatially resolved atlas was necessary. We integrated 259 samples from 201 patients, including distinct biological and treatment backgrounds. Besides, we leveraged the refined cellular populations described here to perform spatial deconvolution uncovering distinct spatial niches that were further validated in a large cohort of bulk samples.

The presence of cytotoxic T cells is essential for antitumor response. Here, we observe that in addition to CD8+ T cells being lower in number (**Figure 1D and Supplementary** Figure 2E), they also exhibit reduced cytotoxic activity in PDAC compared to the adjacent tissue, indicating that the acquisition of the dysfunctional phenotype occurs locally (**Figure 2F**). Later, we discovered that most of these CD8+ T cells enriched in the adjacent tissue are CD8-TRM (**Figure 2B and Supplementary** Figure 3H). These cells exhibited higher expression of activation, cytotoxic, and effector markers in the adjacent tissue (Supplementary Figure 3G). Recent studies uncovered that CD8-TRM cells are essential for immunotherapy response and sustained tumor immunosurveillance (92–94). Thus, the lower number of CD8-TRM cells coupled with their attenuated cytotoxic profile can partially explain the limited effect of neoadjuvant immunotherapy in PDAC (95–98).

The impact of chemotherapy on malignant cells is well documented, however, the effect on CD8 T cells remains elusive. Besides, most clinical trials in PDAC combine chemo with immunotherapy. We did not observe any significant enrichment in T cells from treated patients compared to untreated ones (**Figure 1E**). Differential expression analysis revealed downregulation of tumor reactivity and immunotherapy response genes in CD8-TEX cells from treated patients (Supplementary Figure 3C). This could justify the lack of a robust response in patients undergoing chemoimmunotherapy (95,98). Neoadjuvant chemotherapy was associated with CAFs enrichment, especially iCAF (**Figure 3F-G**). Overexpression of complement genes and CCL19 by CAFs was associated with high T cell infiltration in melanoma (99). However, this was not the case for PDAC. A previous study in breast cancer observed that complement signaling derived from CAFs led to increased recruitment of myeloid-derived suppressor cells promoting T cell dysfunction (100). These findings highlight the crosstalk between iCAF and immune infiltration and unravel a polarization towards iCAF upon treatment.

Our NMF analysis expanded the malignant subtype classification uncovering a subset of malignant cells upregulating MHC-II genes (**Figure 4A-B**). This phenotype was coupled with negative CD4+T and CD8+T cell infiltration and it was enriched in metastasis, including lymph nodes and the liver, indicating a role in tumor progression (**Figure 4C-E and Supplementary** Figure 6B). Moreover, MP2 was positively correlated with the MP5 program, basal/EMT-state, revealing that these cells also show enhanced dissemination capacity (Supplementary Figure 6A). In concordance with our results, a recent study on breast cancer patients and preclinical models showed that metastatic cancer cells in the lymph nodes present upregulation of EMT and MHC-II genes (101). Furthermore, they reported that tumor cells MHC-II+ interact with CD4+T cells leading to T-cell anergy and CD4 Treg accumulation because these tumor cells lack costimulatory molecules. Other groups, however, observed a positive correlation between tumor cells MHC-II+ and immune infiltration in melanoma and lung cancer (102,103). The presence of this phenotype was linked to increased levels of antigen presentation and immune activation (102,103). A study of melanoma and lung cancer patients treated with immunotherapy unraveled the upregulation of MHC-II in tumor cells as a mechanism of acquired resistance to anti-PD1 (104,105). Nonetheless, other reports observed a positive association between immunotherapy response and the presence of tumor cells MHC-II+ (102,105). In summary, these findings indicate a controversial role of MHC-II expression on tumor cells depending on the type of cancer and therapy (106).

Most studies describing distinct TME subtypes of PDAC came from bulk RNA (76,107,108). However, this technology struggles to capture high levels of cellular heterogeneity. Moreover, the lack of cellular resolution hinders us from fully characterizing the intricate cellular interactions. Therefore, we applied unsupervised clustering on cell-type fractions revealing six distinct tumor immunophenotypes (**Figure 5**). This approach allowed us to expand a recent classification uncovering TME subtypes. The upregulation of MHC-II and APC scores on B cells from the Lymphoid-Rich subtype indicates an active role in antigen presentation (Supplementary Figure 8). In addition, germinal center B cells from this subtype exhibited high expression of CD40 (**Figure 5G**). These results contribute to the emerging evidence that B cells can present antigens to dendritic and T-cells within the TLS (109–114). Lymphoid-Rich and Immune-Fibrotic subtypes exhibited higher expression of ligand-receptor pairs associated with antigen presentation, recruitment of T and B cells, and T cell activation in comparison to Immune-Mixed and Myeloid-Rich (**Figure 5D-G and Supplementary** Figure 9C-D). Surprisingly, a direct comparison between Lymphoid-Rich and Immune-Fibrotic revealed high expression of the pair CXCL13-CXCR5. However, the Immune-Fibrotic subtype was not enriched for the TLS signature. A possible explanation is that CXCL13-CXCR5 interaction is not restricted to TLS and has been observed in lymphoid aggregates (114). Moreover, we hypothesize that the Lymphoid-Rich subtype could represent patients in a “late-stage” antitumor response as evidenced by a lower proportion of malignant cells compared to other subtypes. Recently, Shu et al uncovered that CXCL13+ lymphoid aggregates in pretreatment are associated with immunotherapy response in hepatocellular carcinoma (HCC). They also identified a distinct TLS morphology associated with areas of tumor regression suggesting that the maintenance of the mature architecture is antigen-driven (115).

Spatial transcriptomics analysis revealed distinct niches in PDAC (**Figure 6**). Further deconvolution of these niches in bulk data combined with functional gene signatures identified nine clusters. Clusters 2 and 9 were composed mainly of malignant niches with poor immune infiltration and low overall survival resembling desertic phenotypes previously reported (**Figure 7**) (9,43,107). TLS positively correlates with survival in a broad range of tumors, including PDAC, and their location is also relevant for prognosis (114). Most of the TLS found in PDAC are located in peritumoral areas (116–117). Whilst, Cluster 3 exhibited the highest score for TLS niche, effector cells, and Th1 cells signature, this group was also enriched for neutrophils signature and significantly decreased for co-activation and MHC scores (**Figure 7**). All these factors severely hampered the antitumor response in this group. A recent study demonstrated that the presence of TLS in patients with a high neutrophil-to-lymphocyte ratio does not improve survival in urothelial carcinoma (118). Others reported a positive correlation between neutrophil infiltration and peritumoral TLS density in HCC (119). Despite the high score for myCAF-niche, patients from Cluster 8 also showed increased signals of active immune response evidenced by a high score of co-activation, MHC-I, MHC-II, and B cell signature contributing to good prognosis. Depletion of myCAF promoted the establishment of an immunosuppressive TME characterized by low T cell infiltration and high infiltration of myeloid cells leading to poor survival in mouse models (120). *Col1a1* deletion in aSMA+ cells inhibited B and T cell infiltration also in PDAC models (121). Bhattacharjee et al highlighted the tumor-opposing role of myCAF by mechanically restraining tumor spread through type I collagen mechanosignals (122).

Overall, we show that our single-cell atlas containing 259 samples and almost a million cells, integrated with spatial transcriptomics and bulk deconvolution is a powerful resource for the pancreatic cancer field. We uncovered a novel subtype of malignant cells MHC-II+ associated with tumor progression and immune exclusion. Moreover, unsupervised analysis unraveled distinct immunophenotypes with unique cell-cell interactions. By leveraging spatial transcriptomics and bulk RNA sequencing, we uncovered unique clusters and functional gene modules associated with clinical outcomes, underscoring the interplay between TME composition and tumor architecture.

## Conclusion

In conclusion, our integrative multi-omics analysis provides novel insights into the cellular and spatial heterogeneity of the PDAC tumor microenvironment, revealing distinct TME subtypes and spatially organized cellular niches that influence tumor progression, treatment response, and patient survival. By leveraging single-cell RNA sequencing, spatial transcriptomics, and bulk RNA sequencing, we uncovered unique clusters and functional gene modules associated with clinical outcomes, underscoring the interplay between TME composition and tumor architecture. These findings highlight the critical role of the TME in shaping the immunosuppressive and chemoresistant landscape of PDAC. Furthermore, our study demonstrates the value of integrating spatial and transcriptomic data to bridge the gap between molecular insights and clinical translation. This comprehensive approach not only enhances our understanding of PDAC biology but also identifies potential biomarkers and therapeutic targets that could inform personalized treatment strategies, ultimately improving outcomes for this highly aggressive cancer.

## Acknowledgments

The study was funded by PRONON.

## Author contributions

G.F.P.M. developed the study concept with the supervision of E.C.F.C., and C.B. G.F.P.M. performed all bioinformatic analyses with support from M.P.L. and J.F. A.B.O. and S.M.S.M. guided histological annotation. G.F.P.M., E.C.F.C., and C.B. wrote the manuscript, and all authors reviewed and approved the manuscript.

## Code availability

The code will be deposited soon in https://github.com/gabriel-pozo/PDAC_Atlas.

---

**Supplementary Figure 1.**
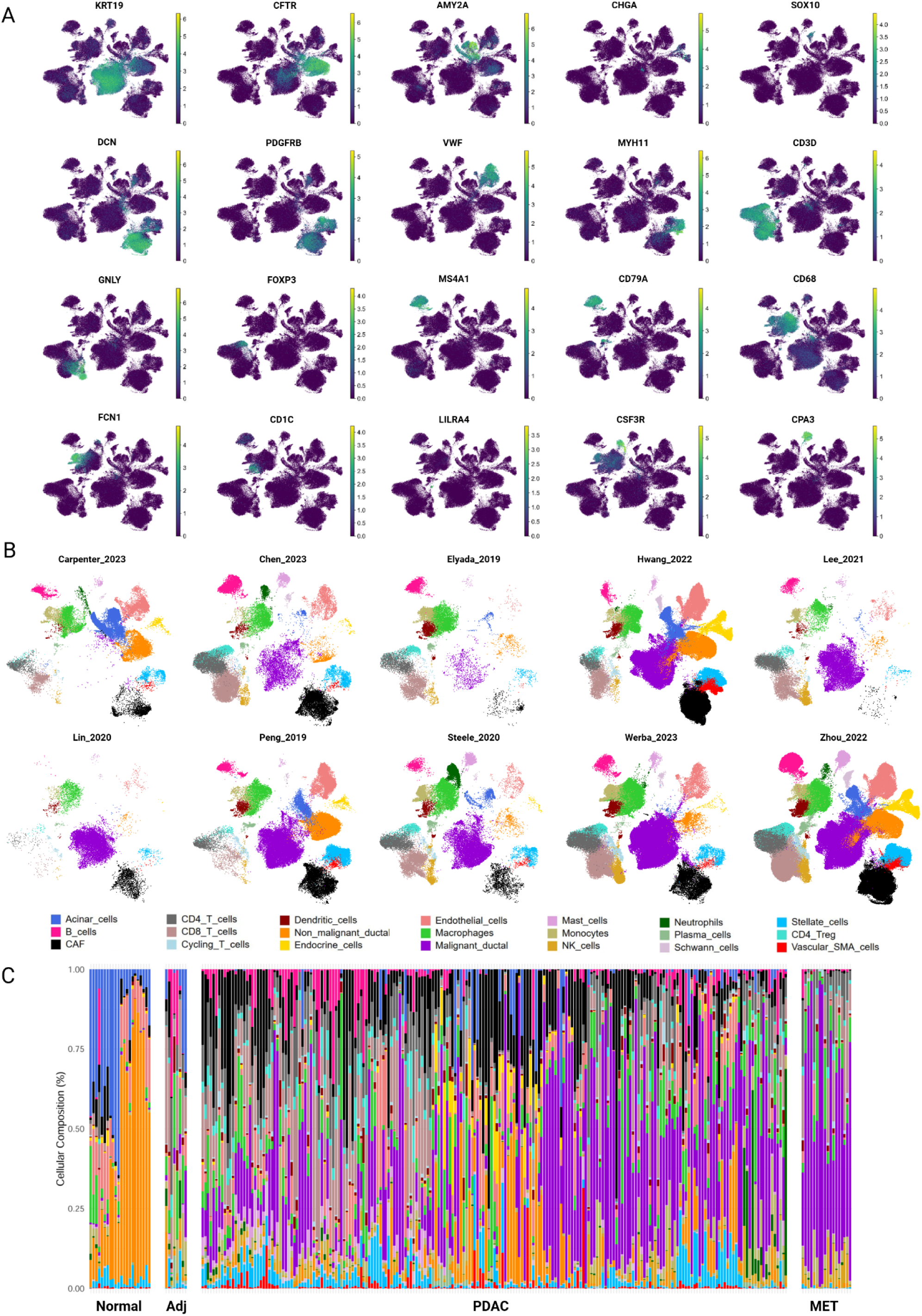
(**A**) UMAP feature plot with the canonical markers used to annotate the major coarse populations; (**B**) UMAP colored by cell type and split by dataset; (**C**) Stacked bar chart for cell type composition by each patient present in the atlas.

**Supplementary Figure 2.**
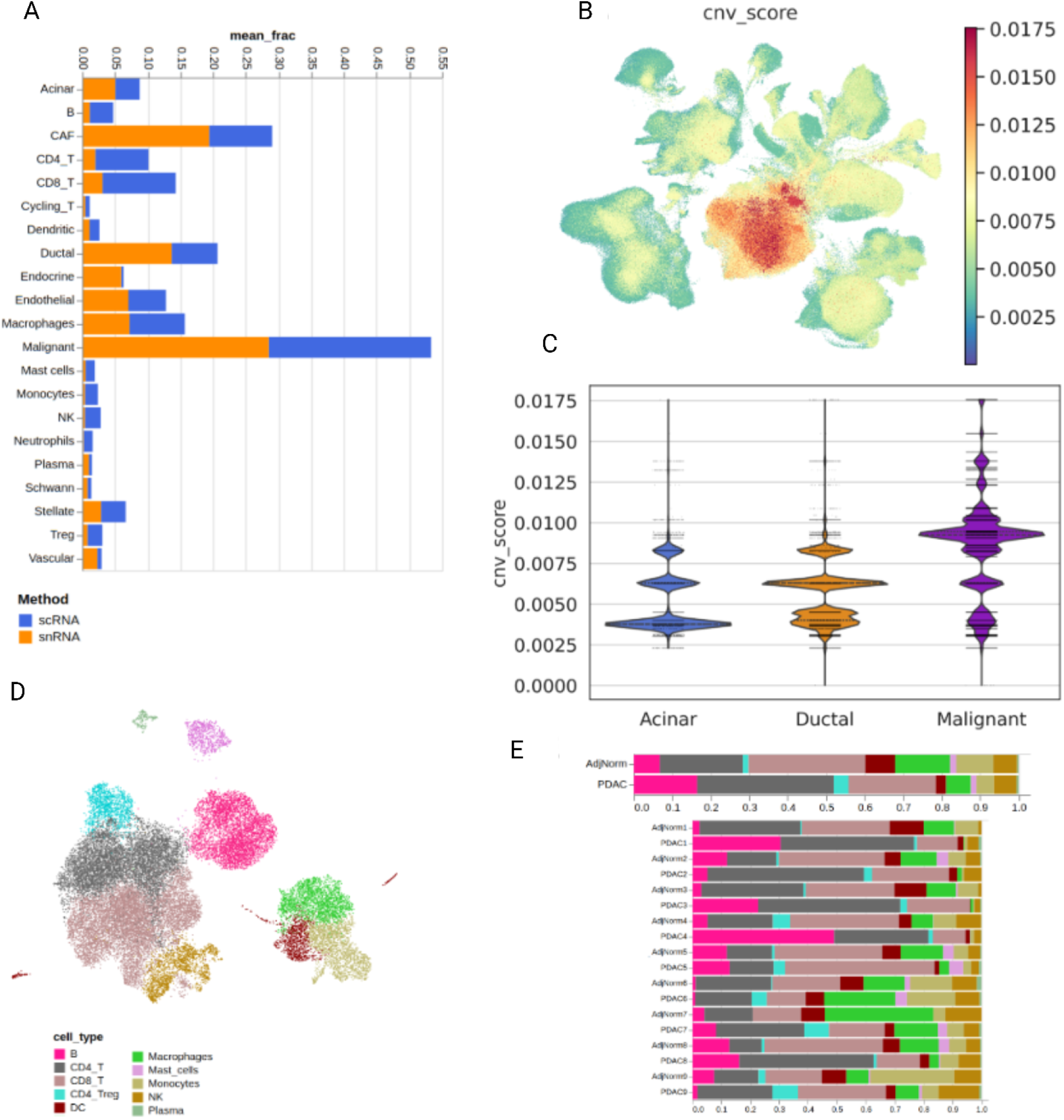
(**A**) Bar chart comparing cell type composition between scRNA and snRNA samples; (**B**) - UMAP feature plot for inferCNV score; (**C**) - Violin plot evaluating inferCNV score across epithelial cells; (**D**) - UMAP projection of Yousuf’s dataset; (**E**) - Top - Stacked bar chart comparing cell type composition between adjacent tissue and PDAC. Bottom - Stacked bar chart by sample.

**Supplementary Figure 3.**
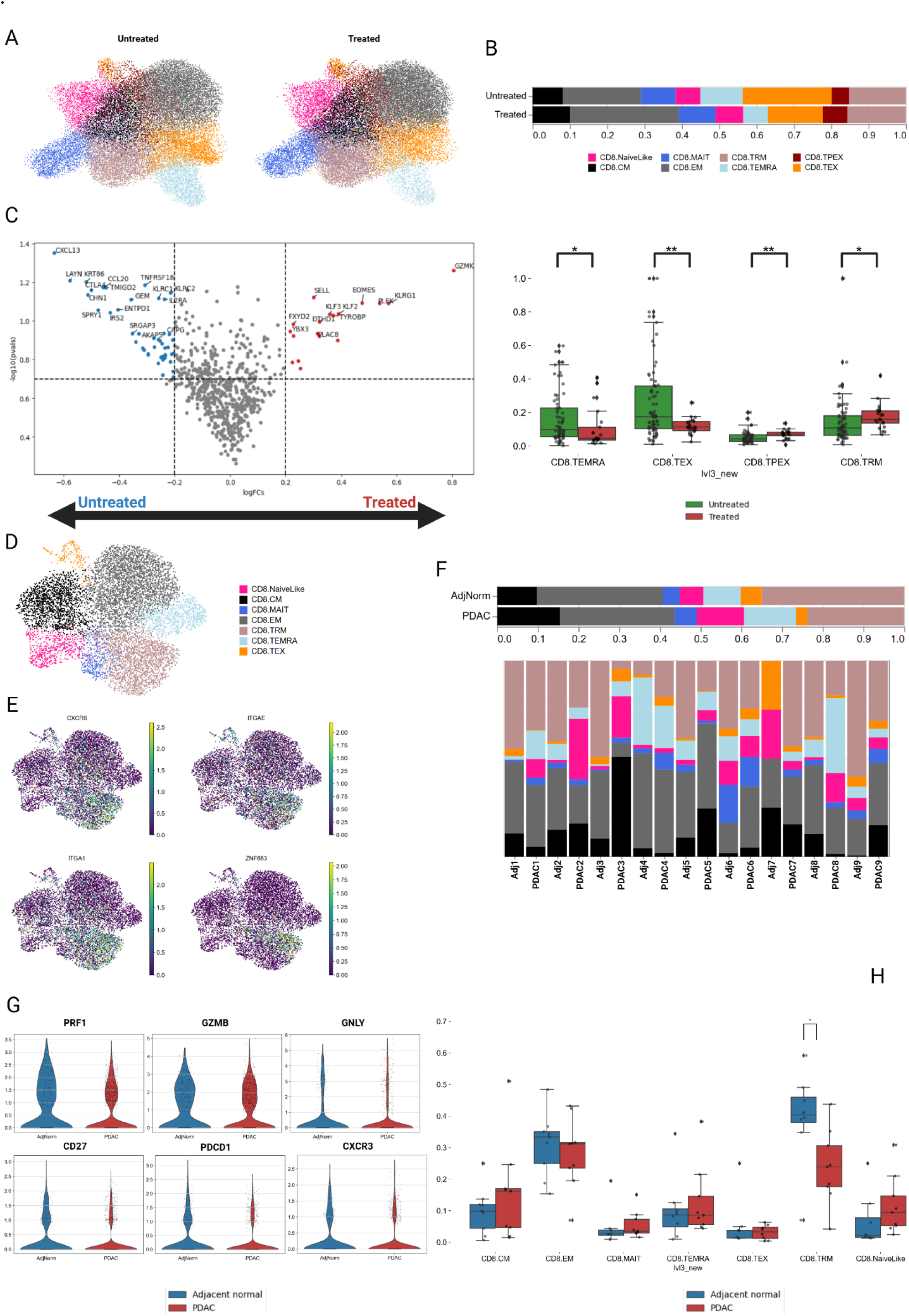
(**A**) - UMAP projection of CD8+T cells split by treatment status; (**B**) - Top - Stacked bar chart for untreated (green) and treated (red) PDAC CD8+T cells composition. Bottom - Wilcoxon test for CD8+T clusters. (**\*** = p < 0.05; **\*\*** = p < 0.01); (**C**) - Volcano plot of differentially expressed genes between untreated CD8.TEX (left) and treated CD8.TEX (right) (FDR < 0.1); (**D**) - UMAP projection of CD8+T cells from Yousuf’s dataset; (**E**) - UMAP feature plot for TRM markers; (**F**) - Top - Stacked bar chart for adjacent tissue and PDAC CD8+T cells composition. Bottom - Stacked bar chart for matched samples CD8+T cells composition; (**G**) - Violin plot evaluating the expression of PRF1, GZMB, GNLY, CD27, PDCD1, and CXCR3 in CD8.TRM from adjacent (left) and PDAC (right); (**H**) - Wilcoxon test for CD8+T clusters between matched adjacent and PDAC. (**\*** = p < 0.05).

**Supplementary Figure 4.**
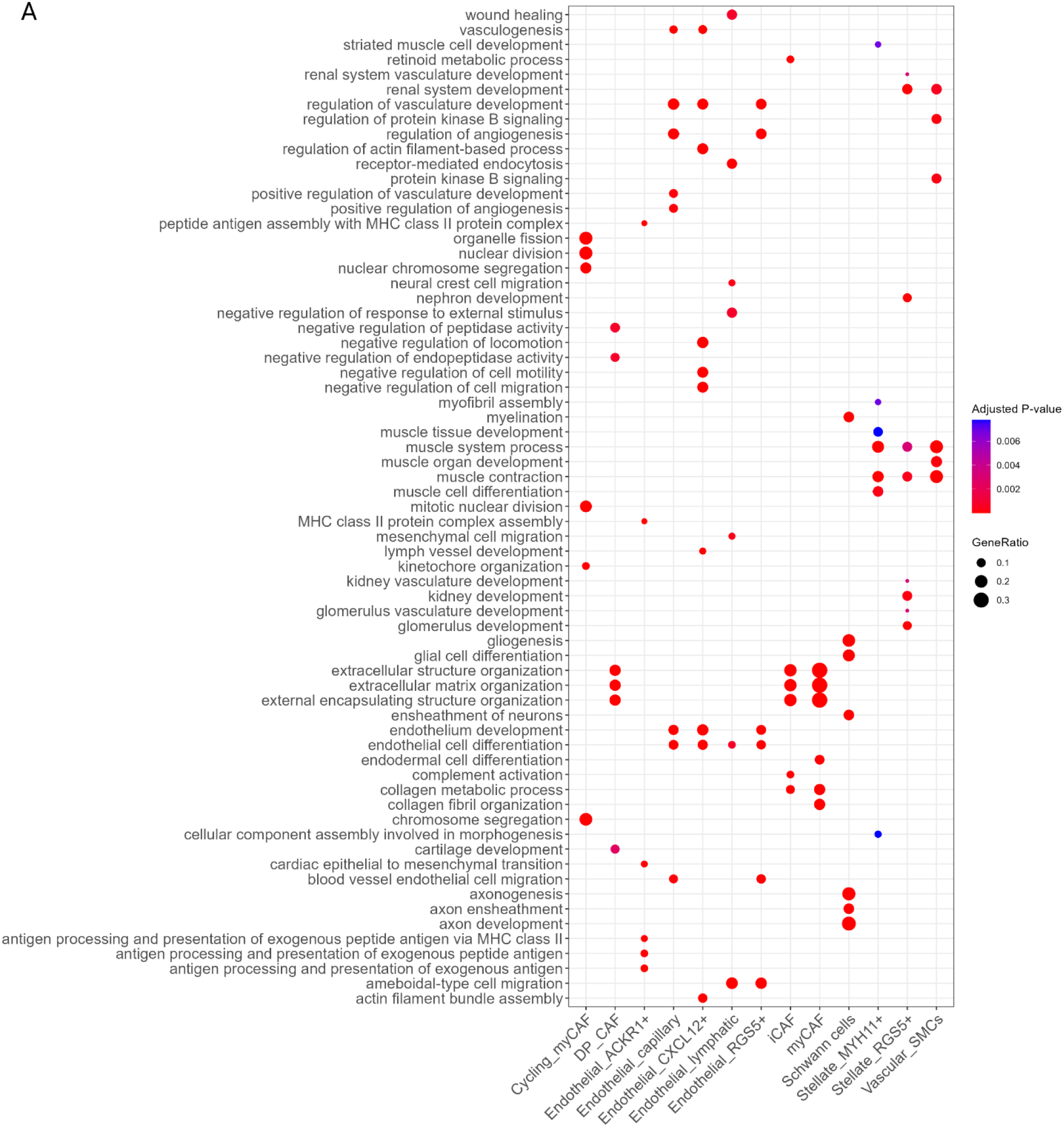
(**A**) - Gene ontology for stromal cell clusters. Rows indicate the pathways obtained from the BP (Biological Process) of GO. Columns represent the stromal cell clusters.

**Supplementary Figure 5.**
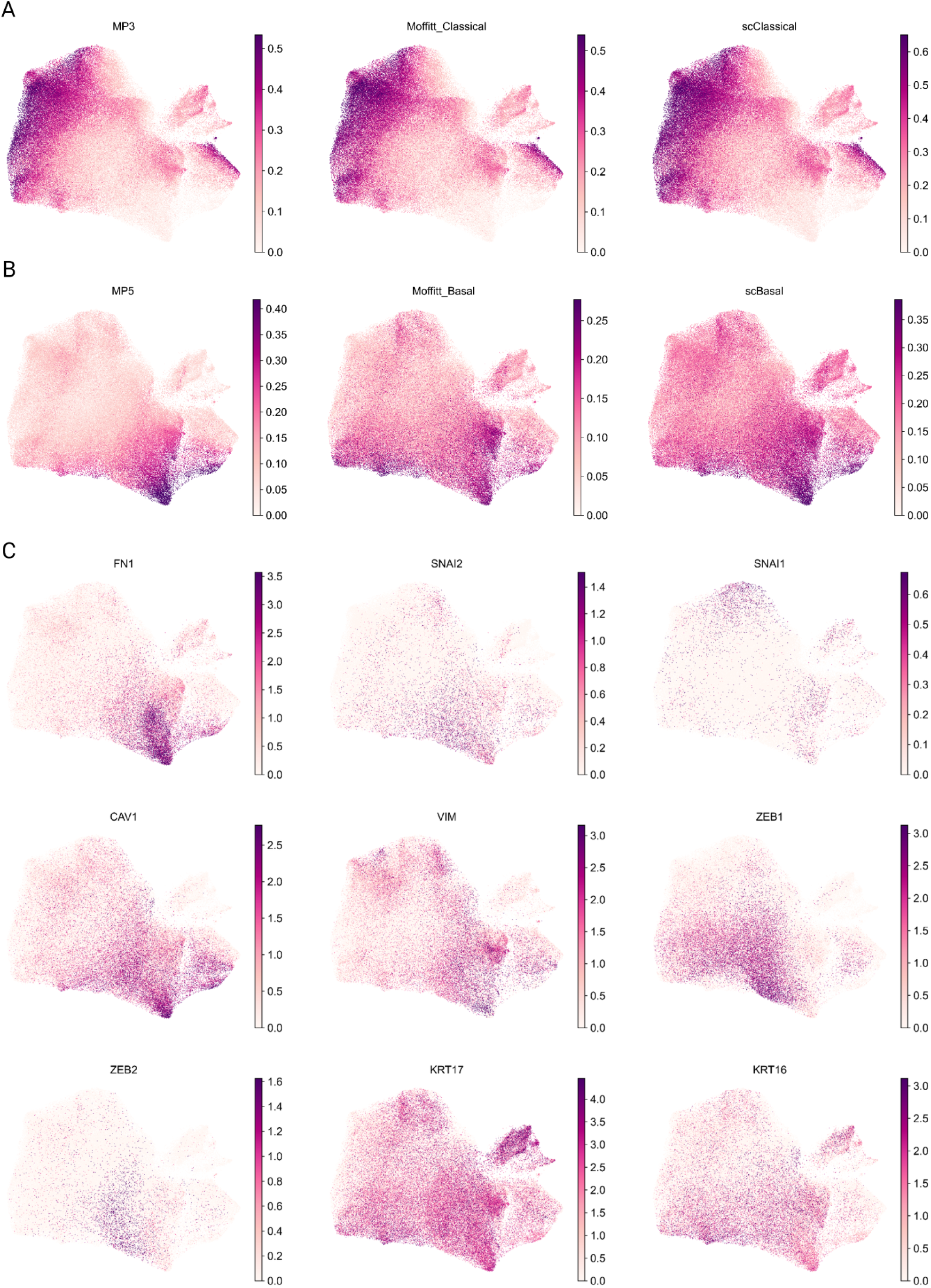
(**A**) - UMAP feature plot for MP3, Moffitt_Classical, and scClassical signature scores; (**B**) - UMAP feature plot for MP5, Moffitt_Basal, and scBasal signature scores; (**C**) - UMAP feature plot for EMT related genes.

**Supplementary Figure 6.**
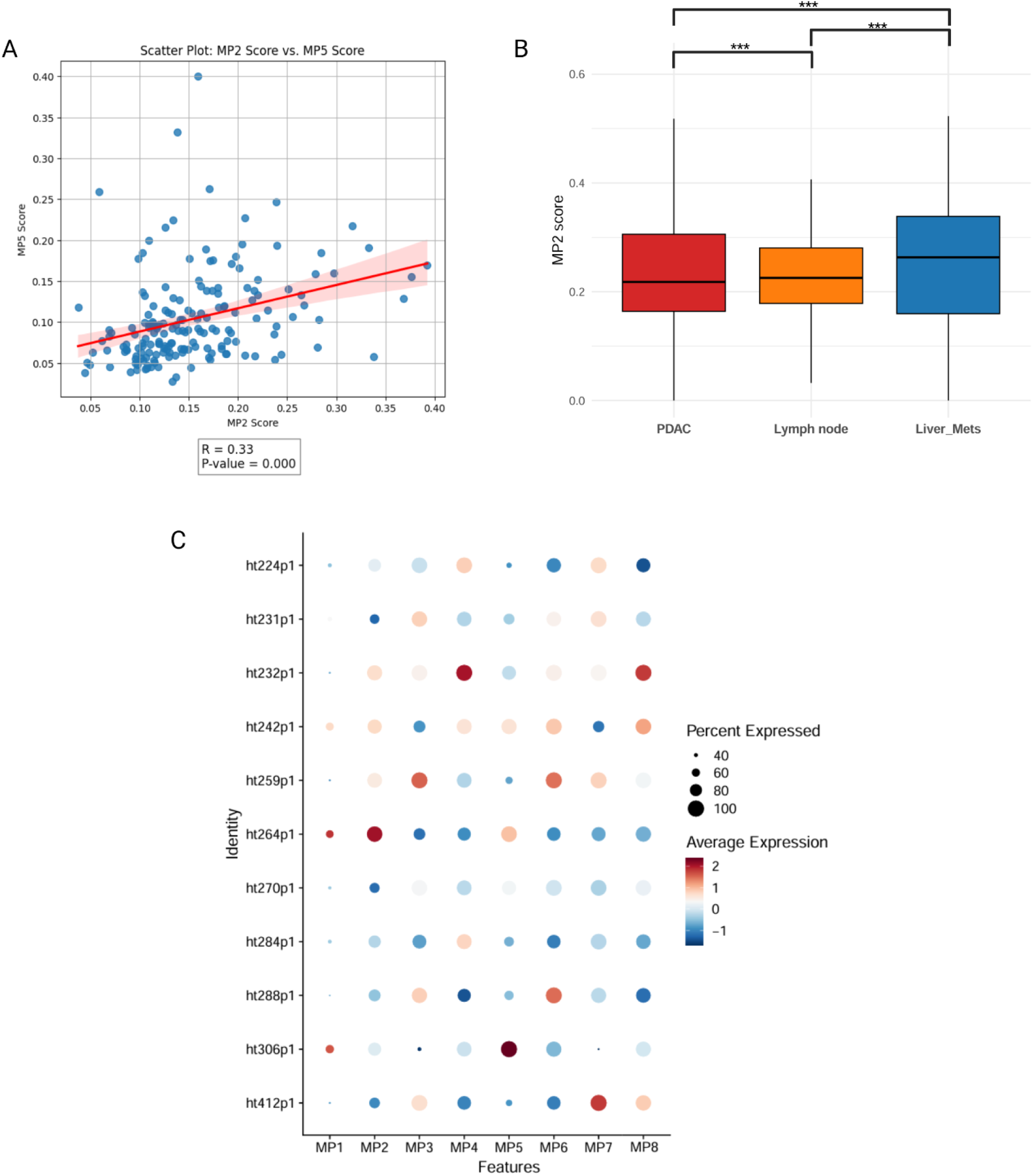
(**A**) - Scatter plot showing the correlation between MP2 score and MP5 score; (**B**) - Box plot for MP2 score across PDAC, metastatic lymph node, and liver metastasis. (Kruskal-Wallis test; **\*\*\*** = p < 0.001); (**C**) - Dot plot for malignant MP’s scores across patients.

**Supplementary Figure 7.**
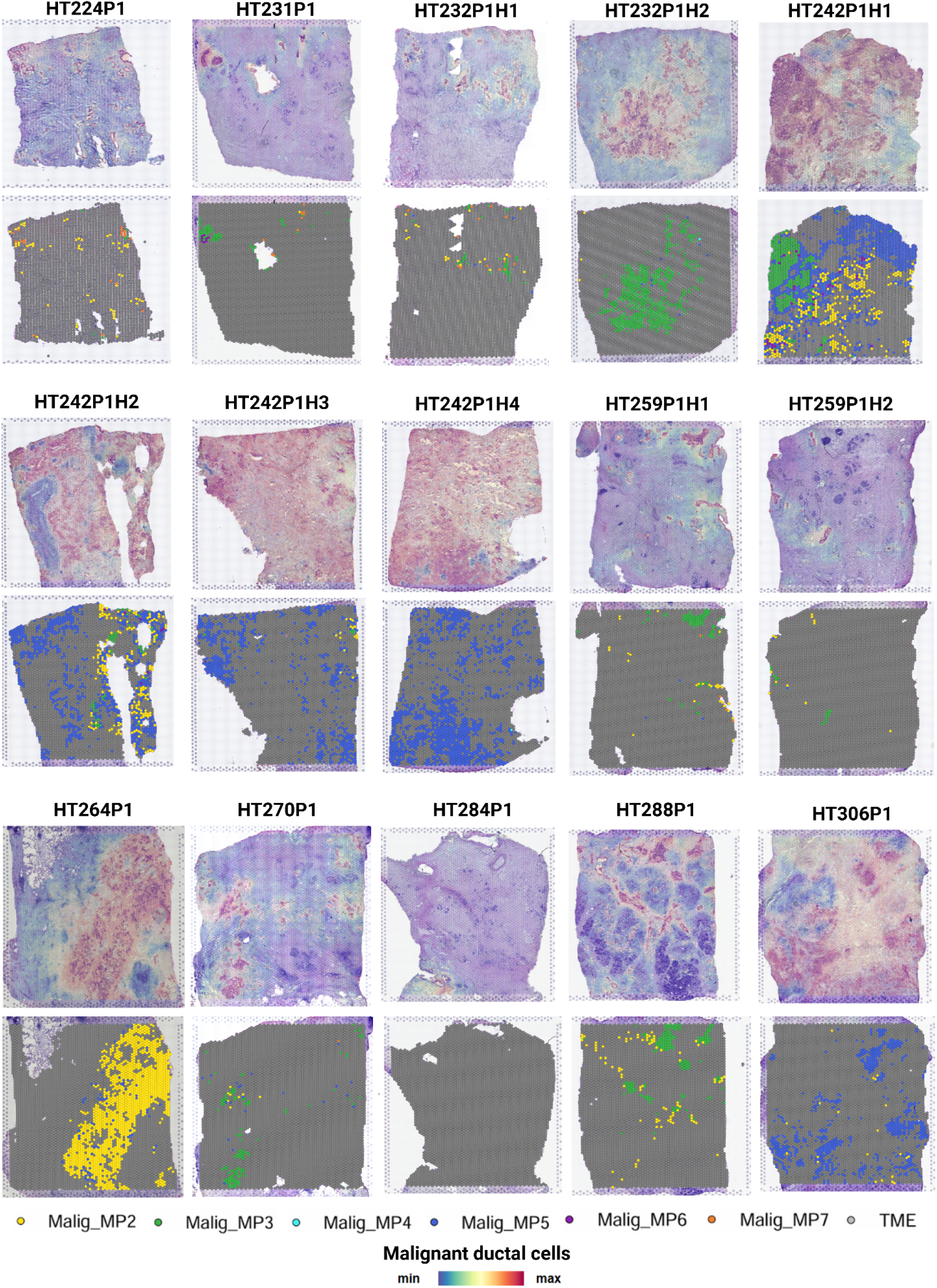
Deconvolved spatial plots showing the proportion of malignant cells and localization of malignant spots assigned to each malignant MP.

**Supplementary Figure 8.**
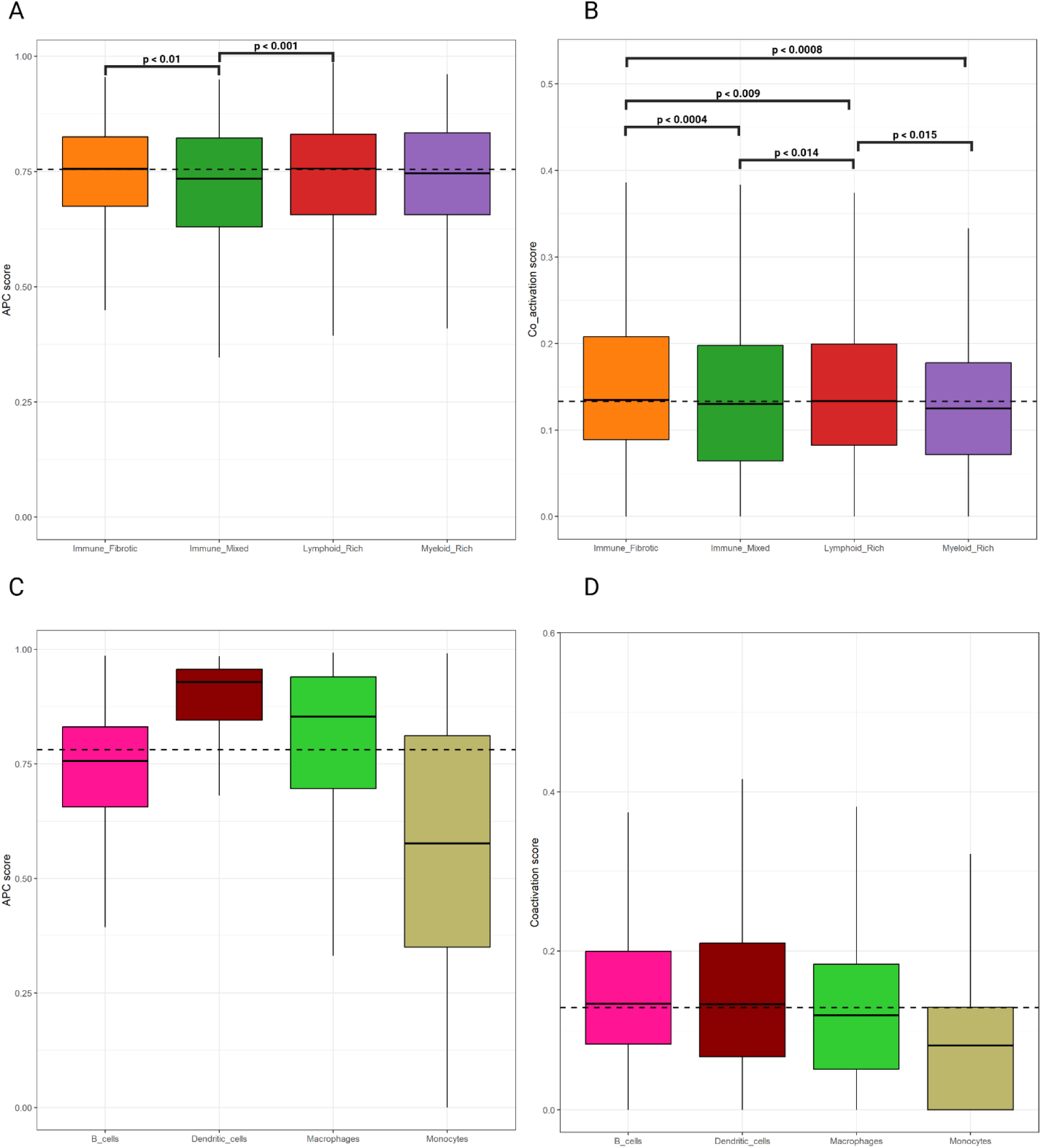
(**A**) - Box plot for coactivation score in B cells among TME subtypes (Wilcoxon test); (**B**) - Box plot for APC score in B cells among TME subtypes (Wilcoxon test); (**C**) - Box plot for APC score among B cells, Dendritic cells, Macrophages and Monocytes from Lymphoid-Rich subtype; (**D**) - Box plot for coactivation score among B cells, Dendritic cells, Macrophages and Monocytes from Lymphoid-Rich subtype.

**Supplementary Figure 9.**
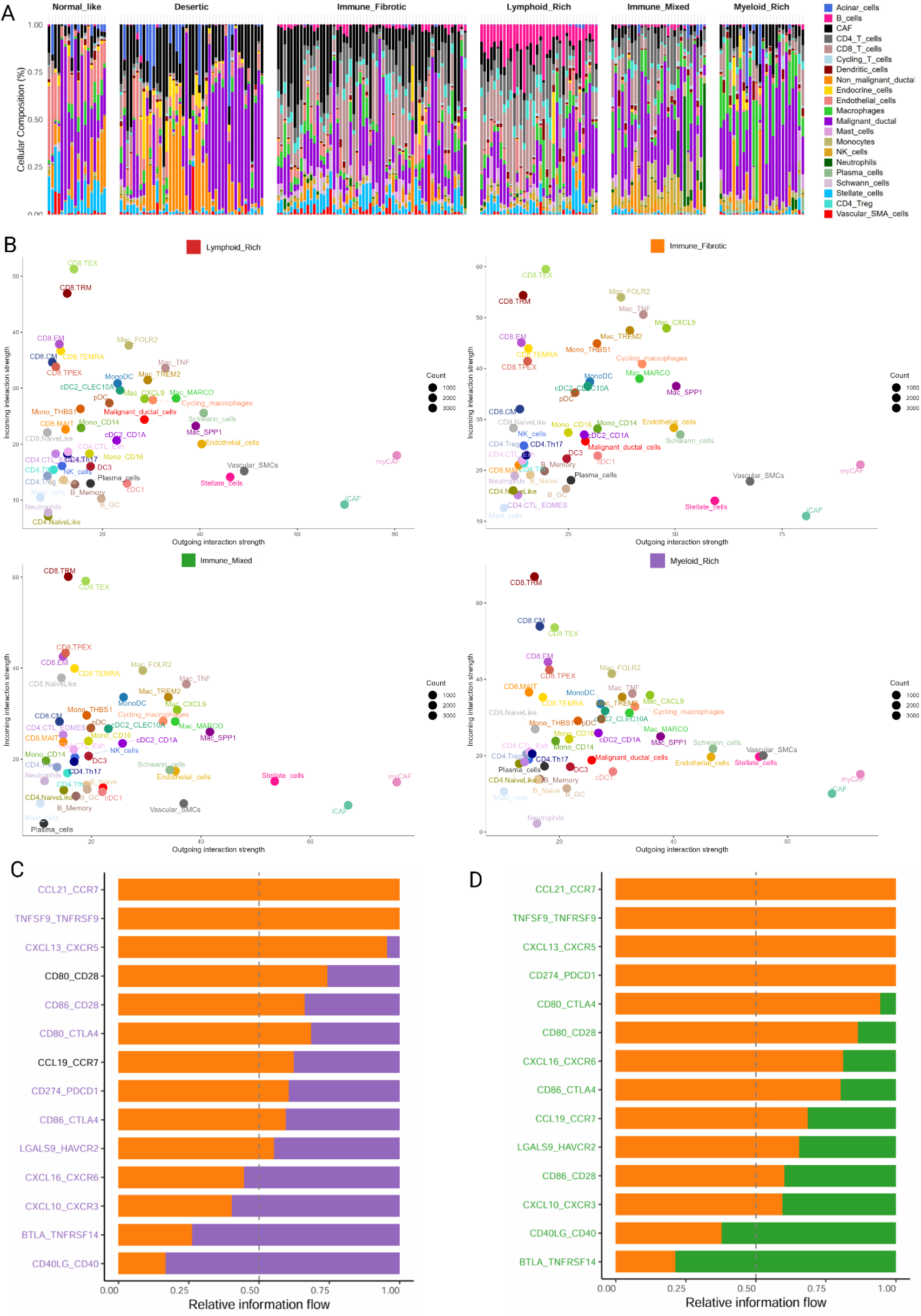
(**A**) - Stacked bar chart for patient cell type composition across TME subtypes; (**B**) - Scatter plot for incoming (y-axis) and outgoing (x-axis) interactions strength of each TME subtype; (**C-D**) - Bar plot for ligand-receptor pair showing the relative flow information.

**Supplementary Figure 10.**
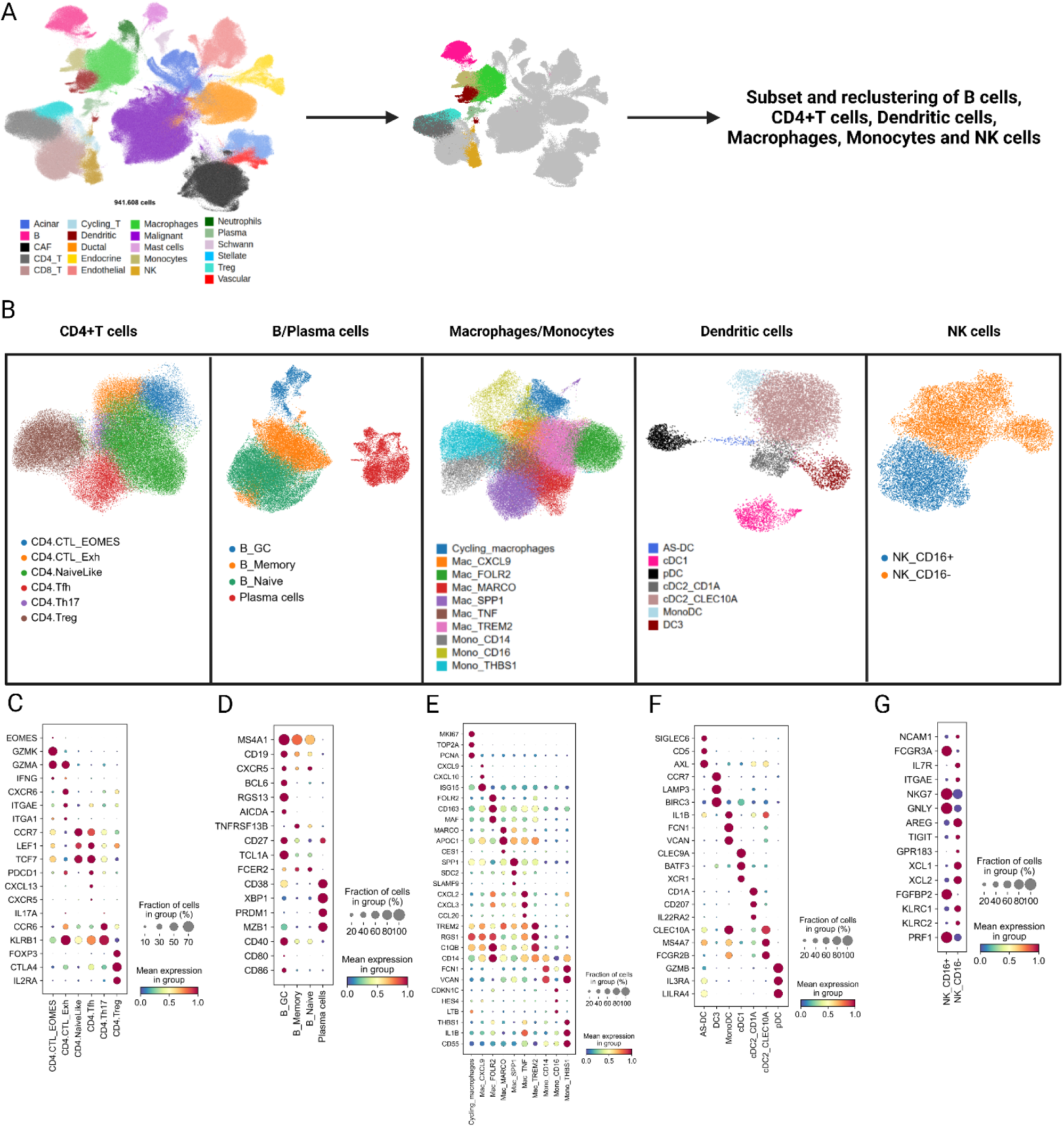
(**A**) - Workflow of the refined cellular annotations for immune cells; (**B**) - UMAP plot showing the refined annotation for CD4+T cells, B cells, Macrophages, Monocytes, Dendritic and NK cells; (**C-G**) - Dot plot showing the main markers used for immune cells characterization.

**Supplementary Figure 11.**
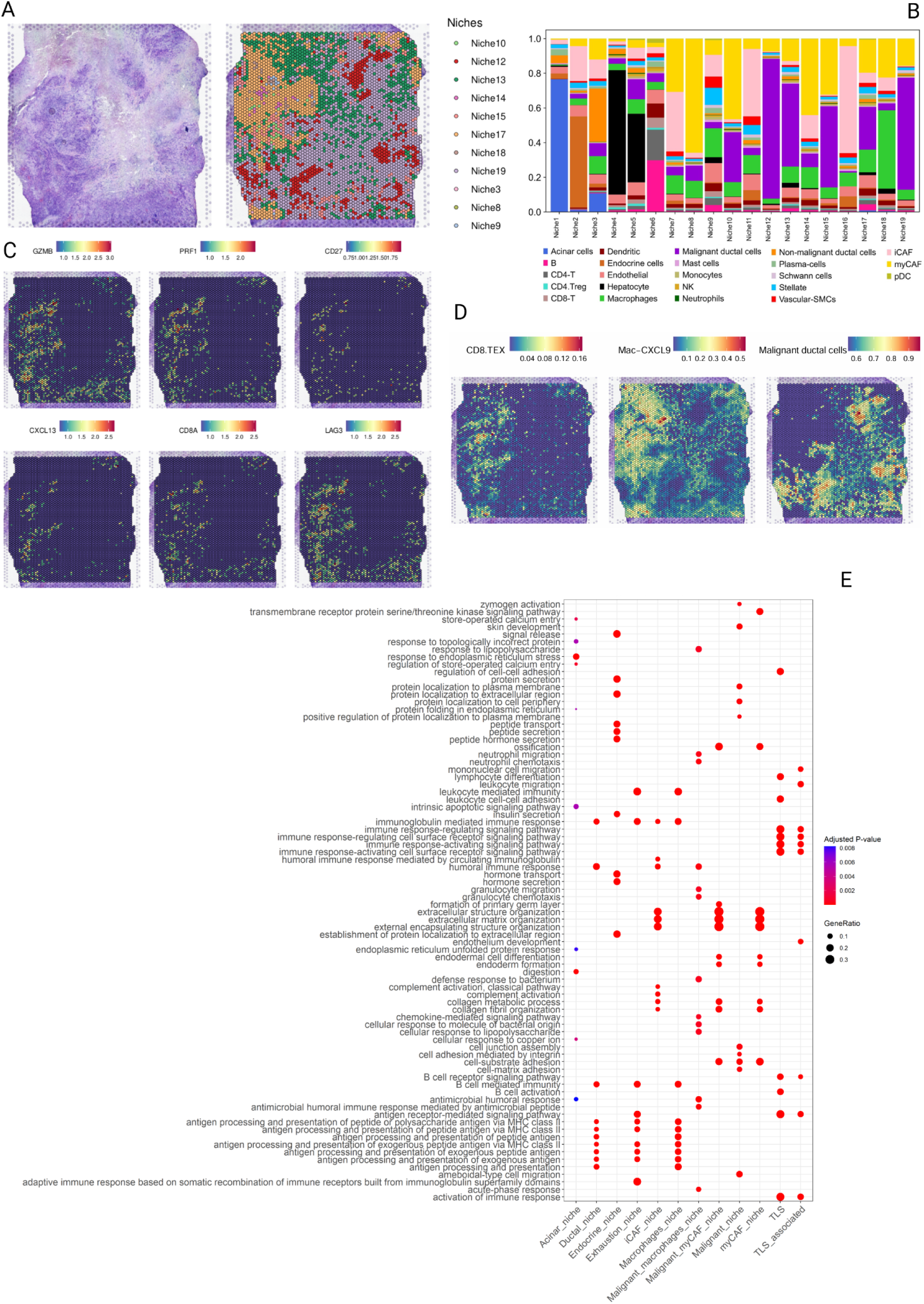
(**A**) - Spatial niche distribution in patient HT306P1; (**B**) - Stacked bar chart showing cell type composition per niche; (**C**) - Spatial feature plot exhibiting expression of GZMB, PRF1, CD27, CXCL13, CD8A, LAG3 in patient HT306P1; (**D**) - Spatial plot showing the co-localization of CD8-TEX and Mac-CXCL9 and malignant cells spatial exclusion; (**E**) - Gene ontology for spatial niches. Rows indicate the pathways obtained from the BP (Biological Process) of GO. Columns represent the spatial niches.

